# Computational and Proteomic Analyses Reveal Cardiac Dysfunction and Heart Failure-Associated Biomarker Secretion from Venezuelan Equine Encephalitis Virus TC83-infected human IPSC-derived cardiomyocytes

**DOI:** 10.64898/2026.04.22.720215

**Authors:** Stephanie V. Trefry, Layan Wahdan, Tyler DiGangi, Caden Andberg, Maame Konadu, Lorreta Opoku, Shannon D Walls, Maria F Galarza, Weidong Zhou, Farhang Alem, Aarthi Narayanan, Qi Wei, Elsa Ronzier

**Author notes:** Equal contribution (Stephanie V. Trefry, Layan Wahdan, Tyler DiGangi).

## Abstract

Arthropod-borne pathogens, many of which are neurotropic, can disseminate beyond the central nervous system to infect peripheral organs. In recent years, an increasing number of cardiac dysfunctions have been reported following arthropod-borne viral infections; however, the mechanism underlying these cardiac manifestations remains poorly understood. In this study, we investigated the impact of Venezuelan Equine Encephalitis Virus (VEEV) TC-83 infection on cardiac function and immune-response of human induced-pluripotent stem cell (hIPSC)-derived cardiomyocytes (hIPSC-CMs). We first confirmed the successful differentiation of hIPSCs into spontaneously beating hIPSC-CMs. We then demonstrated that these cells are highly susceptible to VEEV TC-83 infection, which induced pronounced arrhythmias and complete cessation of beating within 24 hours post-infection. To quantify these functional changes, we developed a segmentation-free computational pipeline that converts frame-to-frame motion in brightfield time-lapse movies into a one-dimensional signal reflecting contractile activity and extracts beat timing, beat rate, and rhythm-regularity features in the time and frequency domains. This analysis revealed progressive disruption of beating dynamics following VEEV TC-83 infection, with early rhythm instability and complete loss of coordinated beating by 24 hours post-infection. Furthermore, mass spectrometry analysis of VEEV TC-83-infected hIPSC-CMs supernatants revealed the presence of biomarkers typically associated with heart failure in patients, underscoring a virus-induced cardiac functional impairment. Together, these findings provide new insight into cardiac complications associated with arthropod-borne viral infections and may support advances in preventive medicine.

## Introduction

Cardiovascular disease (CVD), including coronary heart disease, stroke and heart failure, remains the leading cause of mortality worldwide and represents a growing health burden^1–3^. Increasing evidence demonstrates that viral infections can induce cardiac dysfunctions in heathy individual, and exacerbate pre-existing cardiovascular conditions of heart disease patients ^4–7^. Indeed, CVD patients, known to have dysregulated or impaired immune response, are at elevated risk of developing severe complications after viral infection ^8–10^. Reflecting the bidirectional interaction between infectious diseases and cardiovascular complications, both acute and long-term cardiovascular sequelae have been reported after Influenza ^11^ and SARS-CoV 2 infections ^7,12–14^, as well as chronic cardiovascular impairment associated with persistent viral infections such as HIV ^15–17^ or Hepatitis C virus ^18–21^. These cardiac dysfunctions may arise from direct viral invasion of cardiac tissues or indirectly through virus-induced acute or chronic inflammatory response. This issue becomes even more alarming when considered alongside projections from the American Heart Association, which estimate that by 2035, over 130 million adults, approximately 45.1% of the United States population, will develop cardiovascular diseases ^2^. This expanding vulnerable population is therefore increasingly at risk of adverse cardiovascular outcomes or premature death.

Compounding this issue, the US has recently experienced the re-emergence of infectious diseases caused by arthropod-borne pathogens including West Nile virus (WNV) ^22,23^, Dengue virus (DENV) ^24–26^ and Venezuelan equine encephalitis virus (VEEV) ^23,24^. Global warming is attributed to the spread of these arthropod-borne infections because the habitat ranges of their host species are increasing further north into climates they normally could not survive in, particularly for different species of mosquitoes^27–29^. All these viruses can cause severe neurological complications including encephalitis, meningitis, or stroke. Additionally, those viruses can trigger new cardiovascular conditions such as myocarditis, arrhythmias, and cardiac failures, while worsening pre-existing heart diseases ^30–34^. Indeed, it has been shown that patients that suffer from severe dengue can experience cardiovascular complications ^30,35,36^. A previous study evaluating host response to viral infections has shown that the inflammatory response and oxidative stress from the Dengue viral infection can lead to cardiac and electrophysiological dysfunction ^32^. While other studies have shown that electrophysiology measurements of infected macaques were affected during VEEV infection ^33,34^.

There is a critical shortage of available vaccine candidates for many of the re-emerging pathogens occurring across the globe. This is highlighted by the fact that there is only one FDA-approved vaccine for CHIKV ^37^ including one with restricted recommendation, one FDA-approved vaccine for DENV ^38^ administrated with limited eligibility criteria and no vaccine has been approved for either WNV or VEEV. Importantly, all four viruses are lacking in an FDA small molecule therapeutic option for clinical management. The use of the attenuated strain VEEV TC-83 for vaccine development has resulted in severe adverse reactions and variable protection from VEEV ^39^.

The mechanisms by which viral infections cause cardiac dysfunction are greatly misunderstood. A major limitation is the lack of a physiologically relevant human cardiac model capable of recapitulating cardiac function and activity to enable the study of the impact of infectious diseases. Recently, the use of human Pluripotent-Stem Cells (hIPSC) has emerged as a powerful and relevant model to study diseases including cardiomyopathies ^40–44^.

Here, we aimed to investigate cardiac dysfunctions and immune response of hIPSC-derived cardiomyocytes (hIPSC-CMs) following VEEV TC-83 infection. We first validated the successful differentiation of hIPSCs into mature, spontaneously beating hIPSC-CMs. Next, we demonstrated that these cells are highly susceptible to VEEV TC-83 infection, which induces pronounced arrhythmias and complete cessation of beating within 24 hours post-infection. Importantly, we developed an image-based analytical framework that enables the automated extraction and quantification of cardiomyocyte beating dynamics from time-lapse microscopy recordings. Using this approach, we highlighted infection-related disturbances in cardiac contractility following VEEV TC-83 infection. Furthermore, mass spectrometry analysis of VEEV TC-83-infected hIPSC-CMs supernatants revealed the presence of biomarkers typically associated with heart failure in patients, underscoring a virus-induced cardiac functional impairment. Together, these findings provide new insight into cardiac complications associated with arthropod-borne viral infections and advance diagnostic and precision medicine to improve patient outcomes.

## Material and Methods

### Cells

*Human Induced Pluripotent Stem Cells* (hIPSC) were purchased (healthy control hIPSC line, female, StemCell Technology, Cat. SCTi003-A) and expanded following the manufacturer’s protocol. Cells were cultured as colonies in mTeSR^TM^ Plus media (StemCell Technology, Cat. 100-0279). To maintain a high quality and identity of hIPSCs during expansion, differentiated colonies were removed by aspiration, and the expression of pluripotency markers was confirmed by immunofluorescence and/or western blot.

*Human IPSC-derived cardiomyocytes* (hIPSC-CM) were obtained by following the differentiation and maturation protocol from the manufacturer. After expansion and identity validation, hIPSCs were seeded as a monolayer using mTeSR supplemented with Y-27632 (Dihydrochloride, StemCell Technology, Cat. 72302) into 24-well plates. After reaching 95% confluency, cells were treated with differentiation media A, B, or C for 8 days from the STEMdiff^TM^ Ventricular Cardiomyocytes differentiation and maturation Kit (StemCell Technology, Cat. 05010). After differentiation, hIPSC-CM were cultured in cardiac maintenance media until identity validation and further experiments. The expression of cardiac Troponin was confirmed by immunofluorescence and immunoblot in parallel with the appearance of tissue formation and spontaneous contractility of hIPSC-CMs.

*Vero cells* were purchased from the American Type Culture Collection (ATCC, Cat. CCL-81). Vero cells were cultured with Dulbecco’s Modified Essential Medium (DMEM, Quality Biological, Cat. 112-014-101) supplemented with 1% L-glutamine (Gibco, Cat. 25030-081), 1% NEAA (Gibco, Cat. 11140-050), 10% fetal bovine serum (FBS, ThermoFisher Scientific, Cat. A52568-01) and 1% penicillin/streptomycin (P/S, Corning, Cat. 30-002-Cl).

All cells were cultured at 37°C and 5% CO_2_.

### Virus strains and Virus Amplification and Titration

*Venezuelan equine encephalitis virus TC-83,* vaccine strain stock was generated through electroporation of in vitro–transcribed RNA synthesized from the pTC83 plasmid, as previously described ^45^. The pTC83 plasmid was kindly provided by Ilya Frolov (The University of Alabama at Birmingham), and viral stocks were verified by qRT-PCR prior to experimental use. *West Nile virus* (WNV, MX H 442, Cat. NR-49927), and *Dengue virus type 2* (DENV2, New Guinea C, Cat. NR-84) were obtained from BEI resources and cultured to generate working stocks.

All three virus stocks were amplified in Vero cells in T-150 flasks seeded overnight to achieve 70% confluence. Monolayers were infected at a multiplicity of infection (MOI) of 0.1 and incubated for 1 hour at 37 °C and 5% CO_2_. After incubation, 20 mL of growth media was added. The supernatant was collected and stored at 80 °C. All virus stocks were titrated by plaque assay on Vero cells.

*Viral titrations were determined using plaque assay.* VEEV TC-83 and WNV plaque assays were performed on Vero monolayers. Vero cells were seeded to obtain a ∼90% confluency in 12-well plates the following day. Cells were infected with 10-fold serial dilutions and incubated for an hour. Cells were overlaid with 2 mL of 0.3% agarose in 2X EMEM without phenol red (Thomas Scientific, Cat. 115-073-101) supplemented with 5% FBS, 1 % P/S, 1% sodium pyruvate, and 1% non-essential amino acids (NEAA). Cells were incubated at 37 °C and 5 % CO_2_ for 48 hours (VEEV TC-83) or 72 hours (WNV). After incubation, cells were fixed with 10% formaldehyde for at least 1 hour, the overlay was removed, and cells were stained with 1% crystal violet in 20% ethanol diluted in deionized water. After ∼10 minutes, at room temperature, excess stain was removed under running water, and plaques were counted. *Dengue virus type 2 (DENV)* plaque assays were performed in Vero monolayers. Vero cells were seeded at 3.5 x 10^5^ cells/well in a 6-well plate (in 10% DMEM) and incubated overnight at 37°C and 5% CO_2_. Serially diluted cell culture supernatants or DENV, in DENV dilution media (EMEM supplemented with 2% FBS, 2mM L-glutamine, 2% Sodium Bicarbonate, and 100 U/mL penicillin-streptomycin), were added to the cell monolayer and incubated for 90 minutes with rocking every 15 minutes. Cells were overlaid with a 1:1 mixture of overlay media (2X EMEM supplemented with 5% FBS, 2% NEAA, 200 U/mL P/S, 2% HEPES, and 2mM L-glutamine) and 2% low-melt agarose were added to each well. After a five-day incubation period, a second overlay (1% low-melt agarose with 2% neutral red solution) was added. The following day, the plaques were counted.

### Western Blot

Cells were cultured into 6 or 24-well plates, washed with PBS (VWR, Cat. 02-0119-0500), lysed with RIPA buffer (ThermoScientific, Cat. PI89900) and harvested to determine total protein concentration of samples using BCA Protein Assay Kit (Pierce, Cat. 23227). All samples were normalized to load equal amount of total protein (between 5-15 µg of proteins) Samples were loaded into a Bis-Tris Mini Protein Gel, 4–12%, 1.0–1.5 mm (ThermoFisher Scientific, Cat. NP0321BOX) at 150 Volt for 60 to 90 min using 1X NuPAGE™ MOPS SDS Running Buffer (ThermoFisher Scientific, Cat. NP0001) in a gel tank (ThermoFisher Scientific). After migration, SDS-Page gel was transferred to a PVDF membrane (0.45 µm, ThermoFisher Scientific, Cat. 88518) with a wet system using the NuPAGE™ Transfer Buffer (ThermoFisher Scientific, Cat. NP0006) and a blot module (ThermoFisher Scientific, Cat. B1000). After the transfer, the membrane was blocked for 1 hour at room temperature with a BSA blocking buffer (ThermoFisher Scientific, Cat. 37520), followed by primary antibody incubation (with a dilution 1:3,000) except for GAPDH (dilution 1:10,000) overnight at 4 °C or 2 hours at room temperature in 1% BSA blocking buffer. Primary antibody solutions were washed for 10 min, 3 times with TBST, and secondary antibody-HRP (Cat. 32460 [Goat anti-Rabbit], Cat. 32430 [Goat anti-Mouse]) solutions were added (dilution 1:10,000 in 1% BSA Blocking buffer) for 60 min at room temperature. Membrane was washed 3 times with TBST, and chemiluminescence was detected using SuperSignal™ West Femto Maximum Sensitivity Substrate (ThermoFisher Scientific, Cat. 34094) and the BioRad Chemidoc Imager. Band intensity was analyzed using ImageJ software (NIH; 1.53q).

Mouse heart tissue lysate was prepared using a C3H/HeN mouse heart that was harvested and frozen from a previous experiment ^46^. The heart tissue was placed in blue lysis buffer (2X Novex Tris-Glycine Sample Loading Buffer SDS (ThermoFisher, Cat. LC2676) T-PER Tissue Protein Extraction Reagent (ThermoFisher, Cat. 78510), 0.5M EDTA pH 8.0 (ThermoFisher, Cat. J15684AE), complete Protease Cocktail tablets (Roche, Cat. 11697498001), 0.1M Na3VO4 (Sigma-Aldrich, Cat. 56508), 0.1M NaF (FisherScientific, Cat. S299100), and 1M dithiothreitol (DTT; ThermoFisher, Cat. R0861) and homogenized using a bead homogenizer. The homogenate was centrifuged; supernatant was harvested and used for Western Blot.

### Microscope Imaging

#### Immunofluorescence on fixed cells

Cells were cultured in a 24-well glass-bottom plate (Cellvis, Cat. P24-0-N), either uninfected or infected as indicated, washed once with PBS, fixed with 4% paraformaldehyde (Image-iT™ Fixation/Permeabilization Kit [ThermoFisher Scientific, Cat. A5818101]) for 10 min at room temperature, washed 3 times with PBS, permeabilized for 5 min at room temperature, and washed 3 times with PBS. Cells were then incubated with the BSA Blocking buffer for 60 min at room temperature and incubated with primary antibody solutions at 1:200 for 2 h at room temperature. Cells were washed 3 times with PBS and incubated with the secondary antibody solutions at 1:2500 supplemented with the Hoechst staining reagent at 1:600 (Invitrogen, Cat. 134406B) for 1 h at room temperature, in the dark. Cells were washed 3 times and imaged using an ImageXpress Micro Confocal High-Content Imaging System (Molecular Device).

#### Time-Lapse Imaging on live cells

Time-lapse Imaging was acquired using a Nikon Eclipse Ti inverted microscope equipped with a top stage incubator (Tokai Hit) combined with the Nikon NIS-Elements imaging software. Cells were placed into a pre-equilibrated top stage incubator (37°C and relatively high humidity) for 5 minutes before the recording. Images of the culture were taken every 100ms for 30 to 60 seconds using brightfield.

### Antibodies used for western blot and/or immunofluorescence

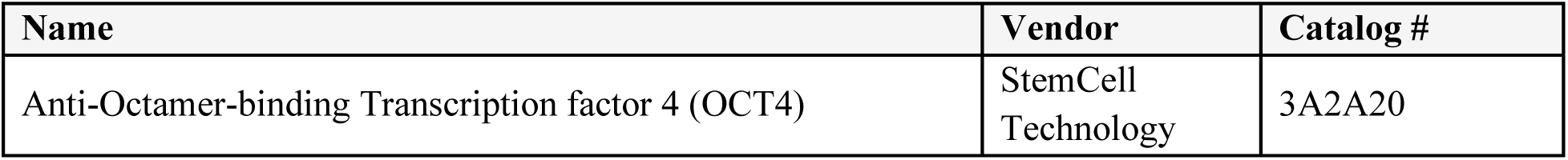
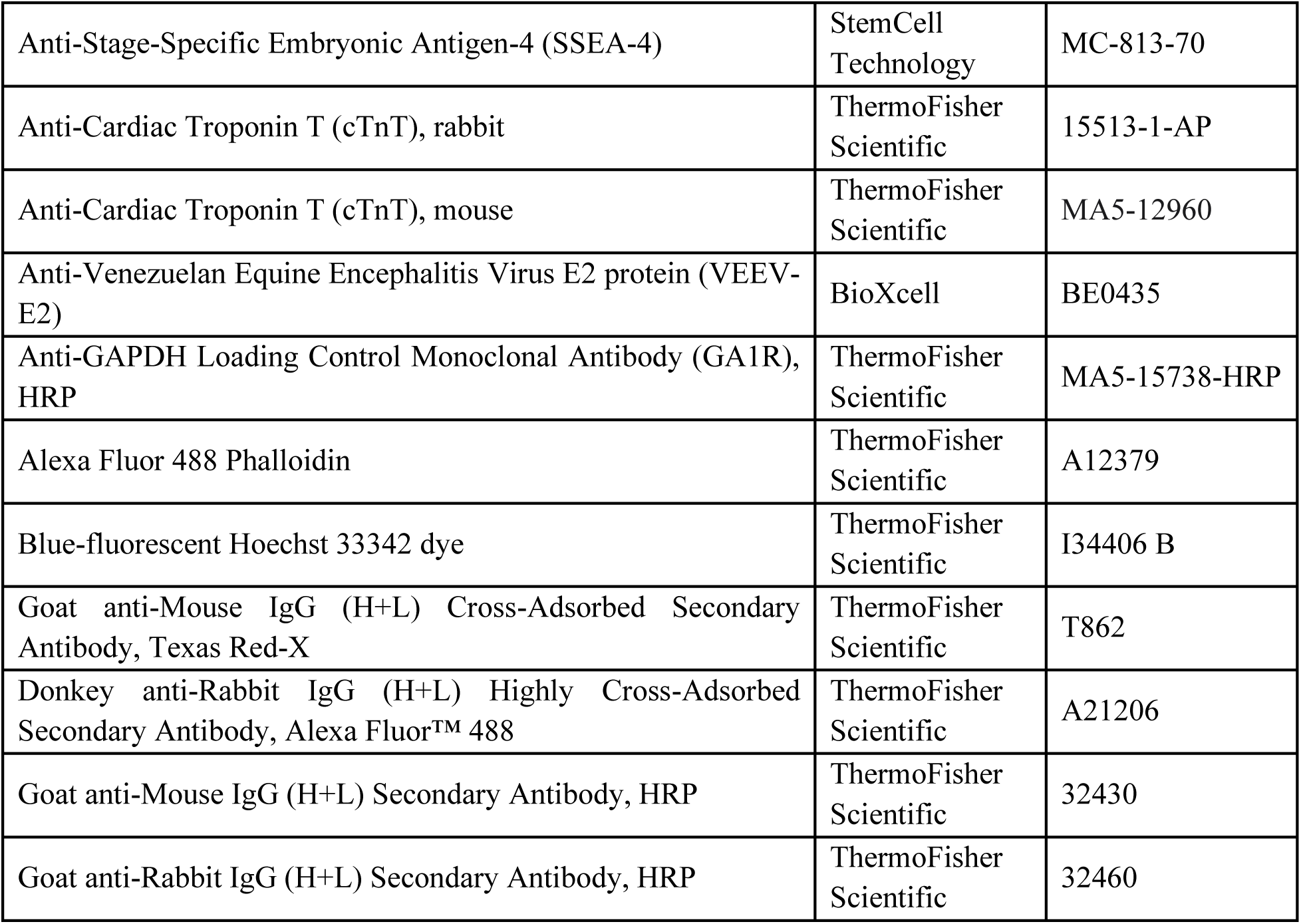

### RNA extraction and quantitative Polymerase Chain Reaction (qRT-PCR)

Extracellular RNA was extracted from cell culture supernatants collected at the indicated time point. Briefly, 50 µL of TRIzol Reagent was added to 50 µL of supernatant, and extracellular RNA was isolated using the Direct-zol RNA MiniPrep kit (Zymo Research, Cat. R2050) following the manufacturer’s instructions. All samples were stored at -80 °C until further analysis. Viral RNA quantification was performed using the StepOnePlus Real-Time PCR System (Applied Biosystems, MA, USA). Viral RNA detection was performed with the Verso 1-Step qRT-PCR ROX kit (Thermo Scientific, Cat. AB4101C) using Integrated DNA Technologies primer pairs (forward: CTGACCTGGAAACTGAGACTATG; reverse: GGCGACTCTAACTCCCTTATTG) and a TaqMan probe (TACGAAGGGCAAGTCGCT-GTTTACC) targeting the nsP1 region of the viral genome. A standard curve was generated using serial dilutions of known VEEV TC-83 RNA concentrations, and absolute viral RNA quantities were calculated in StepOne software v2.3 based on threshold cycle values relative to the standard curve.

### Lactate Dehydrogenase measurements

Level of lactate dehydrogenase was measured with CytoTox 96® Non-Radioactive Cytotoxicity Assay (Promega, Cat. G1781) according to manufacturer’s procedure. To limit the use of samples, 10 µl of samples were used, added to 10 µl of reconstituted subtract mix and incubated for 30 min at 37°C in the dark. After the incubation, 10 µl of stop solution was added and absorbance was measured at 490 nm with a plate reader (Promega).

### Mass spectrometry and analysis

Supernatants from infection studies (mock-treated or VEEV TC-83-infected) were mixed with 20 µL of 8 M urea and incubated at 50 °C for 5 minutes. The mixture was spun at 16,000 g for 2 minutes, and the supernatant was transferred to a clean 0.6 mL tube. The proteins in the supernatant were reduced with 10 mM dithiothreitol, alkylated with 50 mM iodoacetamide, and digested with trypsin at 37 °C for 4 hours. The sample was desalted by ZipTip, dried in SpeedVac, then reconstituted with 10 µL of 0.1% formic acid for mass spectrometry (MS) analysis. Liquid chromatography coupled with tandem mass spectrometry (LC-MS/MS) experiments were performed on an Exploris 480 (ThermoFisher Scientific, Waltham, MA, USA) equipped with a nanospray EASY-nLC 1200 HPLC system. Peptides were separated using a reversed-phase PepMap RSLC 75 μm i.d. × 15 cm long with 2 μm particle size C18 LC column from ThermoFisher Scientific. The mobile phase consisted of 0.1 % aqueous formic acid (mobile phase A) and 0.1 % formic acid in 80% acetonitrile (mobile phase B). After sample injection, the peptides were eluted by using a linear gradient from 5% to 40% B over 90 minutes and ramping to 100% B for an additional 2 minutes. The flow rate was set at 300 nL/min. The Exploris 480 was operated in a data-dependent mode in which one full MS scan (60,000 resolving power) from 300 m/z to 1500 m/z was followed by MS/MS scans in which the most abundant molecular ions were dynamically selected and fragmented by Higher-energy collisional dissociation (HCD) using a collision energy of 27%. “EASY-Internal Calibration”, “Peptide Monoisotopic Precursor Selection” and “Dynamic Exclusion” (15 sec duration), were enabled, as was the charge state dependency so that only peptide precursors with charge states from +2 to +4 were selected and fragmented. Tandem mass spectra were searched against the NCBI human database using Proteome Discover v 2.3 from the ThermoFisher Scientific. The SEQUEST node parameters were set to use full tryptic cleavage constraints with dynamic methionine oxidation. Mass tolerance for precursor ions was 2 ppm, and mass tolerance for fragment ions was 0.02 Da. A 1% false discovery rate (FDR) was used as a cut-off value for reporting peptide spectrum matches (PSM) from the database search.

### Image-based Analytical Framework

A custom computational algorithm was developed in MATLAB (MathWorks) to quantify cardiomyocyte contractile activity from time-lapse brightfield microscopy movies. The algorithm was designed to extract biologically meaningful descriptors of spontaneous cardiomyocyte beating, including contraction timing, rate, and rhythm regularity, without requiring explicit cell segmentation or model-based assumptions. The analysis pipeline consists of video pre-processing, transformation of frame-to-frame motion into a one-dimensional representation of cardiomyocyte contractile activity, and downstream signal analysis in the time and frequency domains.

### Video Pre-processing

Time-lapse movies were imported into MATLAB using the Bio-Formats toolbox to ensure compatibility with microscopy file formats. Each recording consisted of a sequence of brightfield images acquired at a fixed frame rate (100 ms per frame, 10 frames per second) over a duration of 30–60 seconds. Individual frames were converted to grayscale and normalized to ensure consistent intensity of scaling across recordings. Frames were then stacked into a three-dimensional array (x, y, time), enabling efficient frame-wise and temporal processing. No spatial filtering or region-of-interest selection was applied, allowing the analysis to capture global contractile activity across the entire field of view.

### Motion-based Representation of Cardiomyocyte Contractile Activity

To convert image-level motion into a quantitative representation of cardiomyocyte contractile activity, the algorithm computed the **mean absolute difference (MAD)** between consecutive frames. For each time point *t*, the absolute pixel-wise intensity difference between frame *t* and frame *t–1* was calculated, and the resulting values were averaged across all pixels to produce a single scalar value. Repeating this operation across the full duration of the movie generated a one-dimensional temporal signal reflecting global contractile motion over time.

This motion-based signal reflects underlying cardiomyocyte biology: coordinated sarcomere shortening and relaxation produce periodic changes in pixel intensity across the monolayer, while rhythm irregularity, weakened contraction, or asynchronous activity alter the amplitude, timing, and stability of the resulting MAD trace. Because this approach captures aggregate motion rather than individual cell morphology, it is robust to cell shape variability and well suited for detecting changes in collective contractile behavior induced by viral infection.

### Data analysis

The resulting MAD time series was analyzed in both the time and frequency domains to characterize cardiomyocyte beating dynamics. In the time domain, contraction events were identified using peak-detection routines with adaptive thresholding. In cases where baseline drift or amplitude variability limited peak stability, trough-to-trough intervals were used to estimate inter-beat timing. Inter-beat intervals were used to quantify beating rate and temporal variability. To reduce the influence of sporadic noise or abrupt motion artifacts, outlier points exceeding one standard deviation from the median were identified and removed, followed by conditional smoothing using a Savitzky–Golay filter with fixed parameters.

In the frequency domain, Fast Fourier Transform (FFT) analysis was applied to the detrended MAD signal to identify dominant rhythmic components. Prior to FFT, the mean of the signal was subtracted to remove DC offset (constant baseline component), a Hanning window was applied to reduce spectral leakage, and zero-padding was used to preserve frequency resolution. Dominant frequencies and their harmonics were extracted based on peak magnitude, providing a complementary assessment of rhythm stability and the emergence of fragmented or non-dominant frequency components associated with arrhythmic behavior.

### Software availability

The custom MATLAB computational program developed for time-lapse video processing and cardiomyocyte beat analysis will be made publicly available upon publication.

### Statistics

All statistics were calculated from the average of at least three independent experiments using GraphPad Prism (Prism 10 version 10.5.0). Statistics were measured using Welch tests and 2-way Anova tests. The *p* values were indicated as below: * for p ≤ 0.05, ** for p ≤0.01 and *** for p ≤0.001.

## Results

### Characterization of human Induced Pluripotent Stem Cell-derived cardiomyocytes

To generate differentiated human Induced Pluripotent Stem Cells (hIPSC) into mature hIPSC-derived cardiomyocytes, we followed StemCell Technology’s protocol of differentiation (**Figure 1A**). hIPSCs are first expanding as colonies, seeded as isolated cells in the presence of Y-27632, a Rho-associated kinase (ROCK) inhibitor, to prevent cell death, and are cultured until cells reach approximatively 95% confluency. Cardiac differentiation was then initiated using 4 different manufactured STEMdiff cardiomyocytes differentiation basal media: with supplement A for 2 days, with supplement B for 2 days, supplement cardiac C for 4 days. STEMdiff cardiomyocyte maintenance basal media when then used for the remaining time of the experiment (between 3 to 5 days). Representative images of cell organization at each key step are shown in **Figure 1B**.

**Figure 1:**
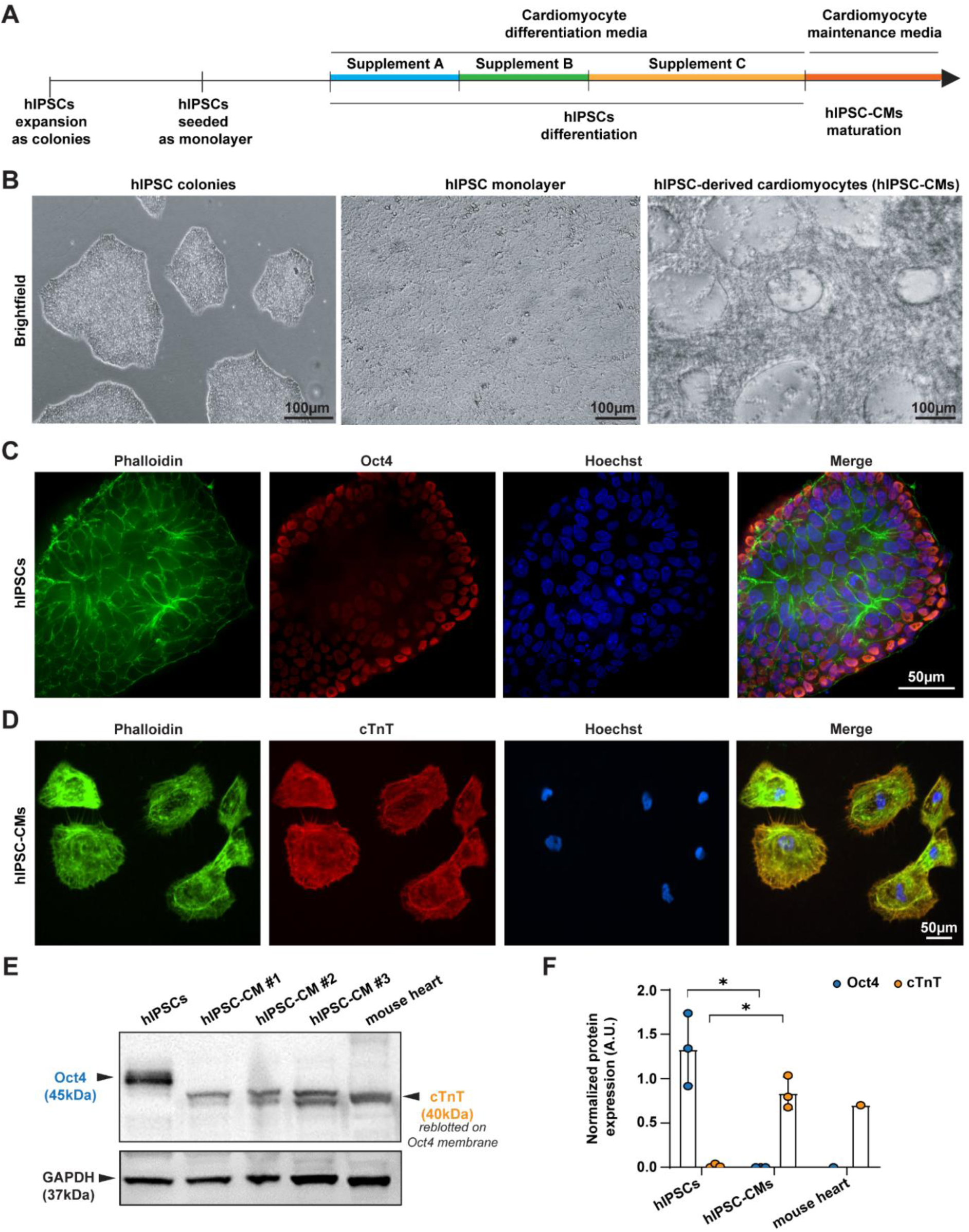
Characterization of Human Induced Pluripotent Stem Cell-derived Cardiomyocytes (hIPSC-CMs). A. STEMdiff ventricular cardiomyocyte differentiation protocol to obtain mature and spontaneously beating hIPSC-derived cardiomyocytes from hIPSCs. B. Representative images of hIPSCs cultivated as colony (left), seeded as monolayer (middle) and hIPSC-CM after differentiation and maturation (right). C Representative fluorescence images of hIPSCs after immunostaining against OCT4, Phalloidin & Hoechst. (D) Representative fluorescence images of hIPSC-CMs after immunostaining against cardiac troponin (cTnT, Ab: MA5-12960), Phalloidin & Hoechst. (E) Representatives Western Blots against OCT4 and cTnT (reblotted on OCT4 membrane) and GAPDH (reblotted on OCT4 and cTnT membrane) using protein lysates from either hIPSCs, hIPSC-CMs (3 independent batch of differentiation), and mouse heart used as a positive control for cTnT expression. (F) Average of normalized Oct4 and cTnT protein expression using GAPDH as a loading control for hIPSCs (n=3 independent batches), hIPSC-CMs (n=3 independent batches) and mouse heart (n=1 heart). Statistics were done using Welch’s t-tests.

To confirm pluripotency of hIPSCs before differentiation, we assessed the expression of Octamer-binding Transcription Factor 4 (OCT4) (**Figure 1C**) and Stage-Specific Embryonic Antigens (SSEA) (**Supplementary Figure 1**) by immunofluorescence, alongside Phalloidin, a cytoskeleton marker and Hoechst, a nucleus marker. A similar characterization was performed to confirm the differentiation of hIPSC into hIPSC-CMs, using cardiac troponin (cTnT), a cardiac marker, alongside Phalloidin and Hoechst staining (**Figure 1D**). Results showed strong expression of OCT4 in hIPSCs and cTnT in hIPSC-CMs (**Figure 1C-D**), consistent with their respective lineage markers. Both OCT4 and cTnT expression were independently confirmed by Western Blot. Protein lysate from a mouse heart was used as a positive control of cTnT expression. (**Figure 1E**). Western Blot quantifications demonstrated a significant expression of OCT4 in hIPSCs compared to hIPSC-CMs and *vice versa*, a significant expression of cTnT in hIPSC-CMs compared to hIPSCs (**Figure 1F**).

Importantly, following this protocol, we reproducibly obtained hIPSC-CMs that exhibited spontaneous contraction with an average beating rate of 43 ± 19 beats per minute (**Supplementary movie 1**). Altogether, those results demonstrated that we have successively differentiated hIPSCs into spontaneously beating, mature, hIPSC-derived cardiomyocytes that can be used for the rest of the study.

### hIPSC-derived cardiomyocytes are highly sensitive to VEEV TC-83

To investigate the susceptibility and impact of viral infection in hIPSC-derived cardiomyocytes, studies were conducted using representative viruses from Alphavirus and Flavivirus genus. First, cardiomyocytes were infected with VEEV TC-83, a live-attenuated vaccine strain derived from Venezuelan equine encephalitis virus IAB at MOI 0.1 either up to 24 or 30 hpi. Supernatants were harvested and cells fixed for immunofluorescence staining. Cells were stained for cardiac troponin (cTnT) to confirm cardiomyocyte lineage and with VEEV E2 protein (VEEV TC-83 E2) to qualitatively assess viral susceptibility. Representative fluorescence images presented in **Figure 2A** demonstrated that hIPSC-CMs are susceptible to VEEV TC-83 infection. The presence of VEEV TC-83 RNA in the supernatant of hIPSC-CM infected cells was confirmed by qRT-PCR with probes targeting nsP1 (**Figure 2B**). The impairment of cell organization of VEEV TC-83-infected hIPSC-CMs MOI 0.1 at 30 hpi compared to mock-treated cells is shown in **Supplementary Figure 2**. To evaluate the capacity of VEEV TC-83 to replicate in hIPSC-CMs, replication kinetics were conducted at 3 different MOIs: 0.1 (low), 1 (moderate), and 5 (high). Supernatants were collected at 0-, 4-, 18-, 24- and 30-hour post-infections. Viral titers were determined via plaque assay. As expected, viral titers at early time points (4 to 18 hpi) increased with MOIs, with MOI 0.1 producing the lower titers, followed by MOI 1 and MOI 5. However, by 24 hpi, all three MOIs reached comparable and high viral titers (**Figure 2C**), characteristic of an alphavirus infection. These results confirmed the infectivity and reproducibility of VEEV TC-83 in hIPSC-CMs.

**Figure 2:**
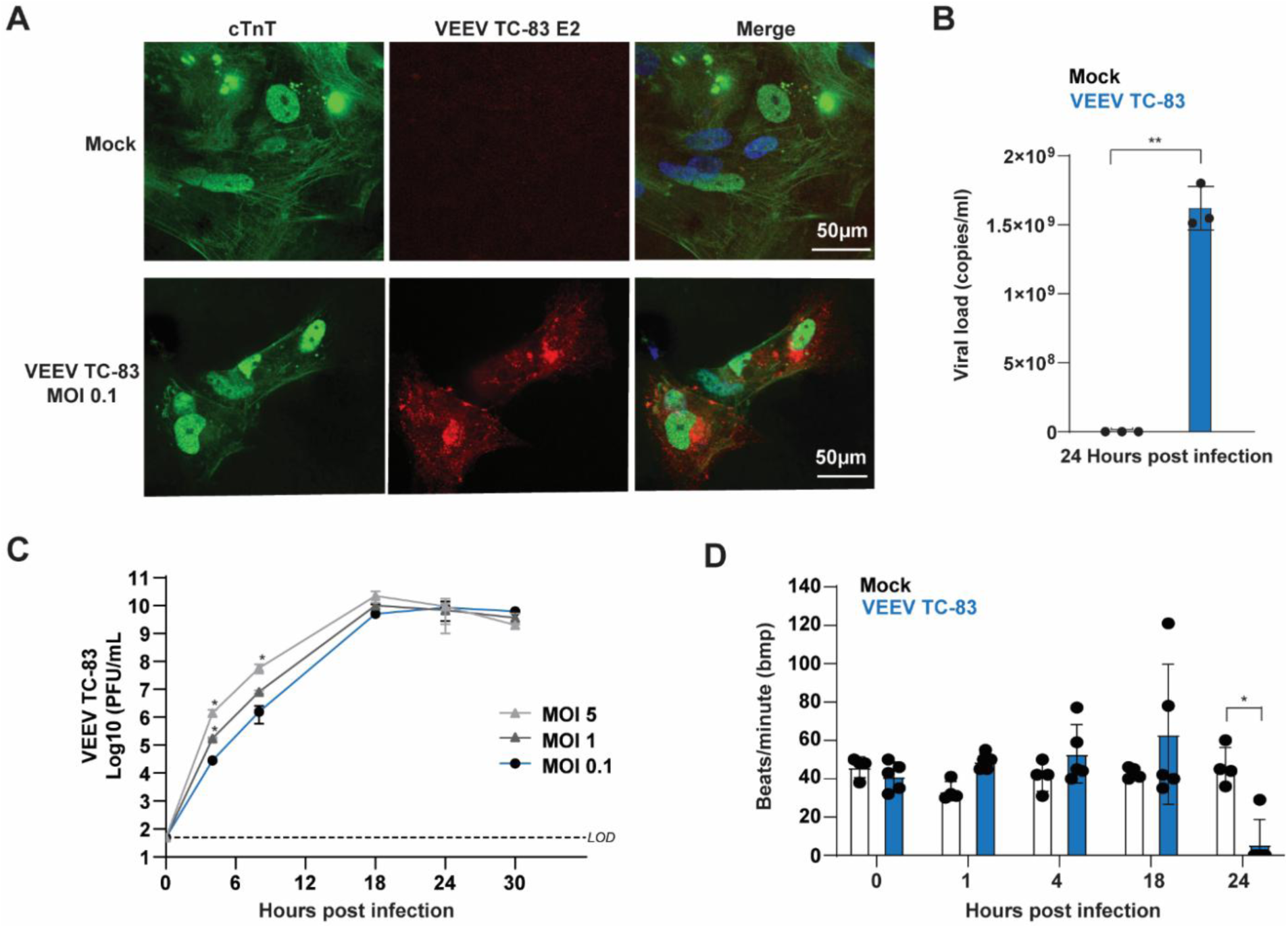
Assessment of VEEV TC-83 virus susceptibility in Human Induced Pluripotent Stem Cell-derived Cardiomyocytes. A. Representative confocal images of hIPSC-CM, either treated as mock or infected with VEEV TC-83 at MOI 0.1 for 24h and immunostained against cardiac troponin T (cTnT, Ab: 15513-1-AP), VEEV-E2 (VEEV TC-83 E2) and Hoechst. respectively. B. Average of VEEV TC-83 viral load (copies/ml) hIPSC-CM either treated as mock or infected with VEEV TC-83 at MOI 0.1 for 24 h. C. Average of VEEV TC-83 titration (PFU/ml) at different MOIs (MOI 0.1, 1 and 5) and at different time points (T0, T4, T18, T24 and T30 hpi). D. Average of beating rhythm (beats/minute) of hIPSC-CMs either treated as mock or infected with VEEV TC-83 after T0, T4, T4, T18, and T24 hpi, * p≤0.05. Statistics were done using Welch’s t-tests.

Using brightfield 60-second time-lapse recordings, we determined, manually, the number of beats per minute (bpm) for both mock-infected and VEEV TC-83-infected cells at different time-points. At early time-points (0 to 4 hpi), both mock-treated and VEEV TC-83-infected cells have comparable beating/rhythm. However, at 18 hpi, we observed arrhythmias, followed by a cessation of beating at 24 hpi in infected cells (**Figure 2D**).

To verify that the observed phenotypes were specifically triggered by a viral infection, we selected two viruses known to induce cardiovascular manifestations and/or myocarditis: Dengue virus subtype 2 ^35,36^ and West Nile virus^31^. We confirmed that both viruses efficiently infected and replicated within hIPSC-CMs, as demonstrated by plaque assay performed at different time points post-infection (Supplementary Figure 3A-B). Consistent with their replication kinetic, viral titers peaked at 48 hpi for both viruses. Using the same strategy as VEEV TC-83, we used brightfield time-lapse recording to determine, manually, the bpm for either mock-treated, DENV-infected or WNV-infected cells at different time points. At 96 and 115 hpi, WNV-infected cells exhibited either a rapid or slow rhythm resulting in large standard deviation across the 4 independent tested wells. However, for DENV-infected hIPSC-CMs, we observed a slowdown at 96 hpi and a significant cessation of beating at 115 hpi (**Supplementary Figure 3 C**).

### Infection of human IPSC-derived cardiomyocytes with VEEV TC-83 induces severe dysfunction revealed by computational image-based analysis

Measuring beating rhythm manually from time-lapse recordings provided useful qualitative insights into cardiomyocyte activity but lacked the resolution needed to characterize beat-to-beat variability, rhythm irregularity, or changes in the frequency content of the motion signal following viral infection. To address these limitations, we developed a computational pipeline to automatically extract and quantify cardiomyocyte beating dynamics from time-lapse microscopy movies. The analysis pipeline processed raw movies through frame extraction and computation of the mean absolute difference (MAD) between consecutive frames, generating a one-dimensional temporal signal that reflects contractile motion over time. All time-lapse movies collected from mock-treated and VEEV TC-83-infected human IPSC-derived cardiomyocytes at multiple time points post-infection were processed using this methodology. The resulting signals were analyzed in both the time and frequency domains to quantify beat timing, beat rate, motion amplitude, and rhythm regularity.

In the time domain, signal peaks corresponding to individual contraction events were identified (**Figure 3**). In the frequency domain, Fast Fourier Transform (FFT) analysis extracted dominant rhythmic components associated with beating frequency (**Supplementary Figure 4**). The algorithm consistently captured beating dynamics in both mock-treated and infected cardiomyocytes at four time points (T0, T4, T18, and T24 hours post-infection). In VEEV TC-83 infected wells, early rhythm irregularity was detectable as early as 4 hours post-infection, progressed to pronounced beat-to-beat instability by 18 hours, and culminated in complete cessation of beating by 24 hours. This cessation of beating was detected in 80 % of the VEEV TC-83 infected wells and was not observed in any mock-treated control wells, which maintained stable rhythmic activity throughout the experiment (**Supplementary Figure 5A-B**).

**Figure 3.**
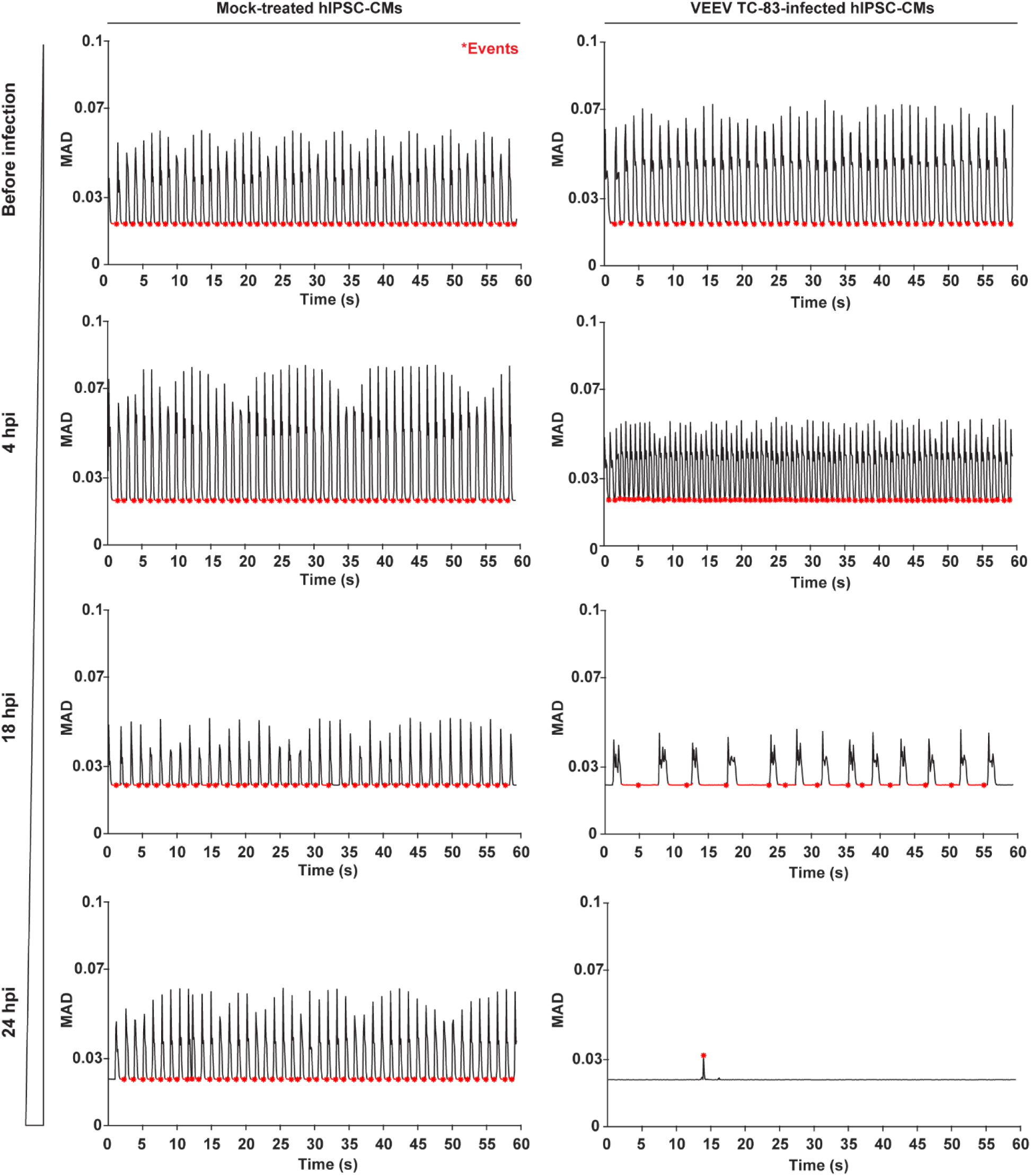
Computational analysis reveals progressive disruption of beating dynamics following VEEV TC-83 infection. Representative MAD time-series traces extracted from 60-second brightfield time-lapse recordings of mock-treated and VEEV TC-83 infected hIPSC-CMs at T0, T4, T18 and T24 hours post-infection. Each trace represents the full-field frame-to-frame motion signal used for time-domain beat detection. Mock-treated cultures show stable periodic oscillations across time, whereas infected cultures show progressive rhythm irregularity and loss of coordinated motion by 24 hpi.

To assess detection accuracy, algorithm-derived beat rate was compared with manually counted beats per minute for each condition and time point. High concordance between manual and computational measurements supported the robustness and reliability of the algorithm for detecting both normal and abnormal cardiac activity (**Supplementary Figure 5C**). Frequency-domain analysis further distinguished infected from control conditions. Mock-treated cardiomyocytes exhibited stable frequency spectra with a consistent dominant peak across all time points, indicative of regular contractile behavior. In contrast, VEEV TC-83–infected cardiomyocytes displayed broadened and fragmented frequency peaks at 4 and 18 hours post-infection, reflecting increased rhythm variability, followed by markedly reduced spectral magnitude and loss of discrete peaks at 24 hours, consistent with beating arrest. Together, these results demonstrate that VEEV TC-83 infection induces progressive and severe disruptions in cardiomyocyte beating rate and rhythm.

Overall, in agreement with qualitative observations from time-lapse movies, computational analysis demonstrated that hIPSC-derived cardiomyocytes undergo pronounced rhythm instability and loss of coordinated contractions following VEEV TC-83 infection, whereas mock-treated controls maintain stable rhythmic activity. These findings establish that the developed computational approach provides a sensitive, non-invasive, and quantitative framework for detecting infection-related disturbances in beating dynamics and support its utility for assessing viral cardiotoxicity and therapeutic protection in future studies. Together, time-domain and frequency-domain analyses provide a quantitative framework linking image-derived motion signals to biologically relevant features of cardiomyocyte function, including beating rate, rhythm regularity, and progression toward arrhythmic or quiescent states.

### Infection of human IPSC-derived cardiomyocytes with VEEV TC-83 triggers a robust host-immune response and the secretion of heart failure biomarkers

We observed a cessation of beating of hIPSC-CMs at 24 hours post-infection with VEEV TC-83. To determine whether this loss of contractility resulted from cellular dysfunction or cell death, we first measured the level of secreted lactate dehydrogenase (LDH), a biomarker of cell death ^47,48^, in mock-treated or VEEV TC-83 infected hIPSC-CMs at different MOI and at different time points (**Figure 4**). We observed no significant difference in secreted LDH level between conditions prior to 30 hpi. At 30 hpi, however, LDH secretion was significantly higher in both MOI 1 and MOI 5 conditions compared to mock-treated cells. These results suggest that the cessation of beating observed in Figure 2D and Figure 3, at MOI 0.1, 24 hpi, is attributed to a cellular dysfunction rather than cell death at this time point.

**Figure 4.**
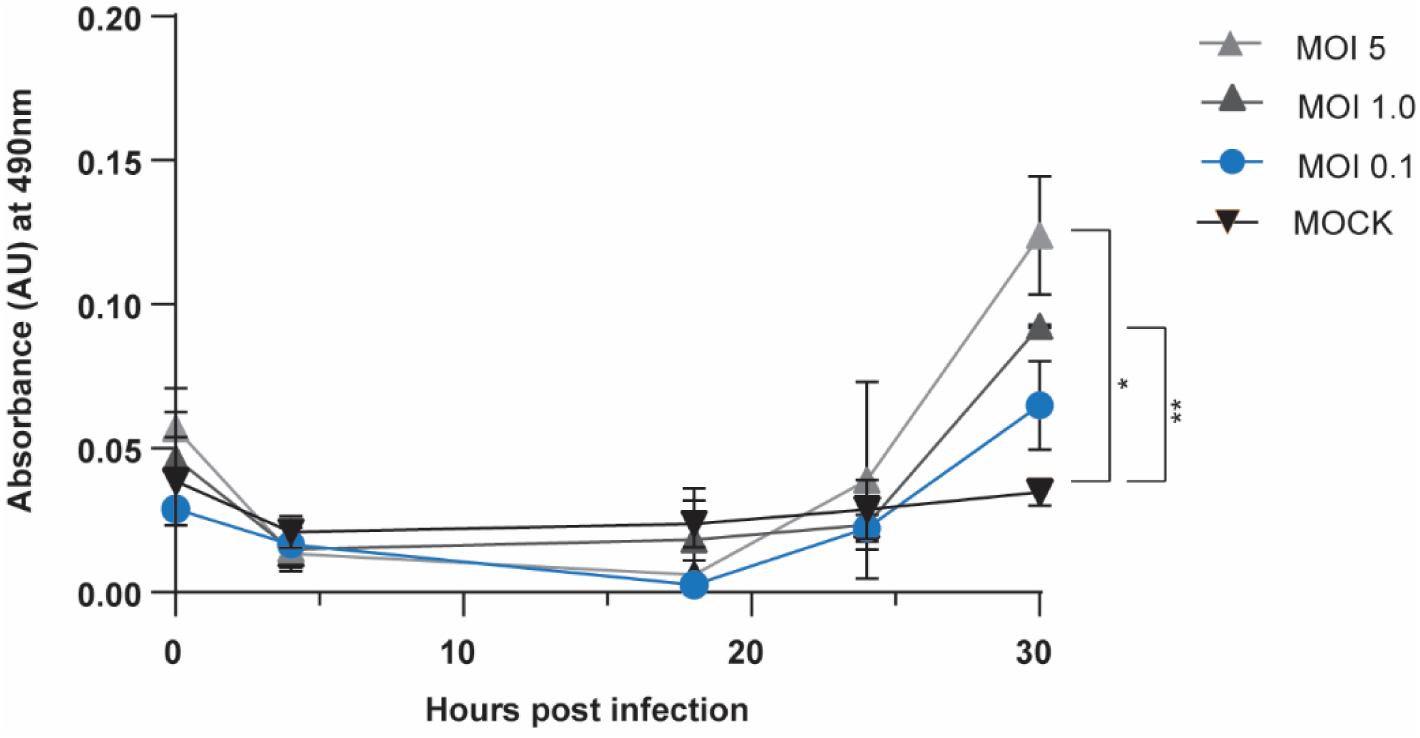
Assessment of Lactate Dehydrogenase (LDH) secretion by hIPSC-CMs following VEEV TC-83-infection. Average of LDH secreted in the supernatant of mock-treated or VEEV TC-83 infected hIPSC-CMs at T0, T4, T18, T24 and T30 hours post-infection. Statistics were done using two-way ANOVA.

To investigate the host-immune response of hIPSC-CMs following VEEV TC-83 infection, we then performed LC-MS/MS-based proteomic analysis of secreted factors at 24 hpi. This approach allowed us to characterize the host response and identify biomarkers indicative of cellular stress and adaptation pathways or necroptotic processes. From this study, we identified 613 unique proteins of which 360 overlapped between infected and uninfected samples, 64 were unique to uninfected samples, and 189 were specific to VEEV TC-83 infection (**Figure 5A**). Using *STRING*-based protein-protein interaction network analysis for the 189 unique proteins identified in the infection condition, we identified multiple functionally relevant protein clusters involved in innate immune systems, signal transduction, infectious disease pathways, metabolic regulation, and cellular homeostasis (**Figure 5B-C**). Among, the secreted proteins identified in the supernatant of VEEV TC-83-infected hIPSC-CMs annotated in Supplementary Table 1, we identified several proteins involved in cardiomyocytes stress response that modulate inflammatory response and reduce damage and/or promote repair such as Macrophage Migration Inhibitory Factor (MIF) ^49^, Glucose-Regulated Protein 78 (GRP78) ^50^, Aldolase A (ALDOA) ^51^, Stress-induced-phosphoprotein 1 (STIP1) ^52^, Protein disulfide-isomerase A3 (PDIA3) ^51^, Cofilin-1 ^35^, and pyruvate kinase (PKM) ^36^. It has been shown that the upregulation of Prostaglandin E2, identified here, affects mitochondria function and reduces contractility in mouse ^54^. Interestingly, several additional identified proteins are known to be secreted in the context of heart diseases. Examples include Phosphoglycerate mutase 2 (PGAM2) which is elevated in serum of patients with tachycardia-induced heart failure ^57^; Nuclear Autoantigenic Sperm Protein (NASP) which promotes cardiac regeneration after myocardial infarction ^56^; the 60 kDa heat shock protein (HSP60) which is highly expressed in the myocardium of patients with heart failure ; Complement factor I is activated in myocardial infarction ^39^; Multidrug Resistance associated Protein 5 (MRP5) which is known to increase in patients with ischemic cardiomyopathies ^60^ and is upregulated in the left ventricular myocardium of dogs with chronic heart failure ^61^; and High Mobility Group Box 1 (HMGB1) which is secreted during ischemia-reperfusion injury of the heart ^62,63^.

**Figure 5:**
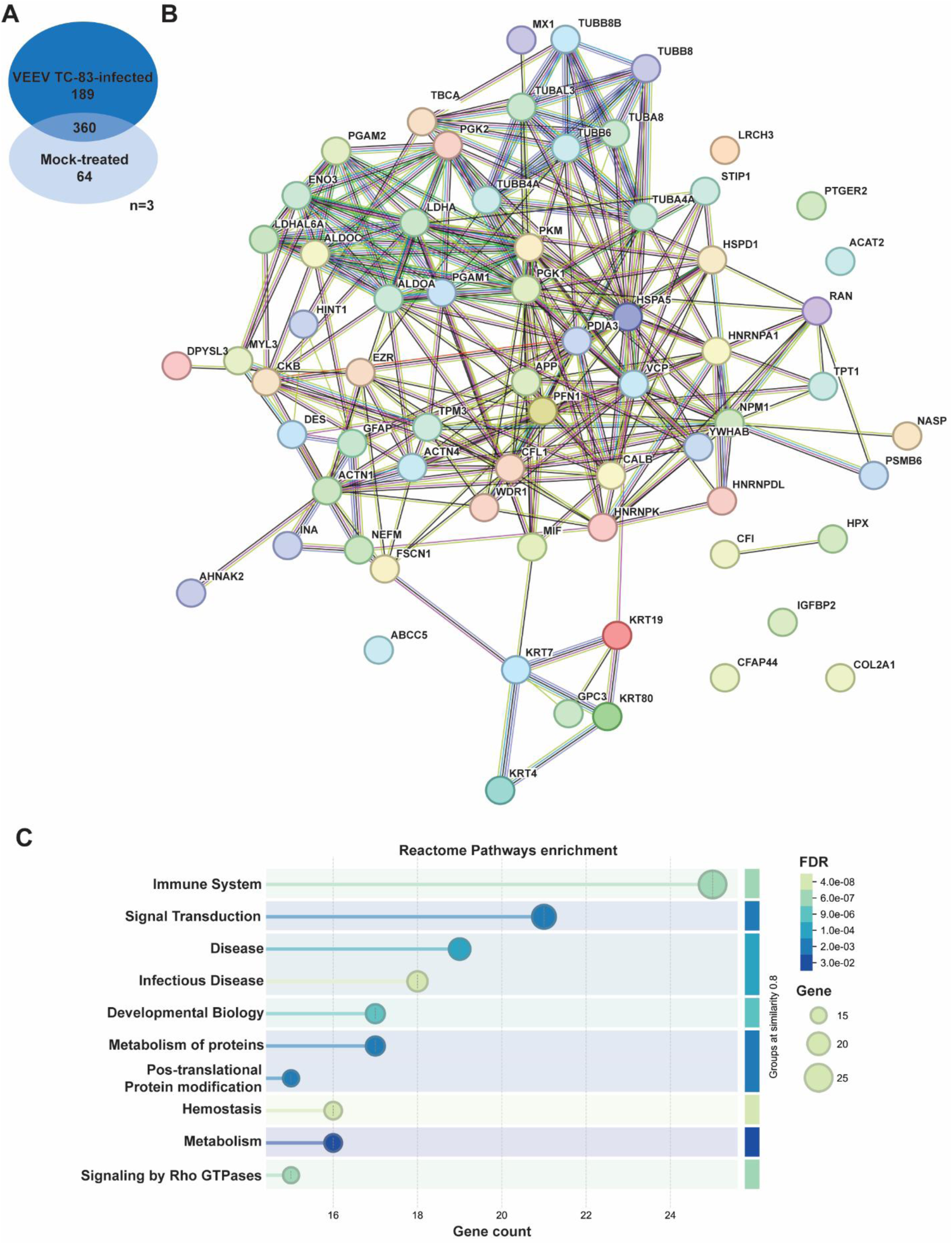
Analysis of secreted proteins from VEEV TC-83 infected hIPSC-cardiomyocytes. hIPSC-cardiomyocytes were infected with VEEV TC-83 and supernatants collected for LC-MS/MS to identify secreted proteins. A. Comparison of identified proteins between VEEV TC-83-infected and mock-treated hIPSC-CM samples. B. STRING analysis of protein-protein interactions identified specific to VEEV TC-83 infected samples. C. Reactome pathway enrichment analysis of upregulated genes from VEEV TC-83 infected samples. Bubble plot showing enriched Reactome Pathways (RP) terms for significantly upregulated genes at 24 hours post infection. Circle size corresponds to the number of genes within each RP category, and color reflects enrichment significance (false discovery rate (FDR) value).

Among the identified proteins, the majority are known to be associated with cardiac hypertrophy and/or contractility dysfunction such as Desmin ^62^, Myosin Light chain 3 ^65^, Tropomyosin a-3 ^66^, Alpha-actinin 1 & 4 ^65^, Profilin ^68^, cofilin-1 ^54^, fascin, ezrin, glycolytic enzymes (PKM, PGAM, ALDOA), and ER/mitochondria stress proteins (GRP78, HSP60, STIP1). This dysfunction may lead to an early cessation of beating preceding cardiomyocytes death, as indicated by an increase level of both Lactate Dehydrogenase ^69^ and HMGB1^62,63^.

Analysis of the VEEV TC-83 hIPSC-CM supernatant secretome revealed biomarkers associated with cardiac inflammation, stress responses, heart diseases, and tissue repair, suggesting that cardiomyocytes actively engage host-defense mechanism for survival against the viral infection before cardiomyocyte death.

## Discussion

In this study, we first confirmed the successful differentiation hIPSCs into hIPSC-derived CMs by validating the expression cardiac Troponin T, a well-established lineage marker ^70,71^, alongside the loss of OCT4 expression, a pluripotency marker ^72–74^, and the presence of spontaneous beating activity ^74^. Using mass spectrometry data, we also identified proteins secreted that can be expressed by cardiomyocytes confirming the lineage of the hIPSC-CMs used in this study. For example, we identified Secreted Protein Acidic and Rich in Cysteine (SPARC), a protein playing a role in cardiomyocyte contractility ^75^; Follistatin-like 1 (Fstl1) is secreted by the heart ^76^; Apolipoprotein E (ApoE) is also secreted by cardiomyocytes^77^.

Following validation of hIPSC-CM lineage, we demonstrated that these cells are highly susceptible to VEEV TC-83 infection. Viral replications reach an average of 1.24e10 PFU/ml at 18 hours post infection even at low MOI, demonstrating extremely high viral production within a rapid kinetic. Functionally, we initially observed manually pronounced arrhythmias and complete cessation of beating within 24 hours post-infection. Those results are coherent with the literature, which shows severe cardiac dysfunctions have been observed after Eastern equine encephalitis virus after 24 hours post infection ^33^.

We further developed a segmentation-free computational pipeline that qualified cardiomyocyte beating dynamics directly from time-lapse microscopy recordings, thereby identifying highlighted infection-related disturbances in cardiac contractility following VEEV TC-83 infection. Importantly, using mass spectrometry analysis of VEEV TC-83-infected hIPSC-CMs supernatants identified the release of biomarkers typically associated with heart failure in patients including PGAM2, NASP, HSP60, CFI, and MRP5, underscoring a virus-induced cardiac functional impairment. Together, these findings demonstrated that cardiomyocytes are susceptible to VEEV TC-83, West Nile and Dengue viruses and bring new insight into cardiac complications associated with arthropod-borne viral infections. This study may support advances in preventive medicine for both healthy individuals and patients with underlying heart disease.

Robust human-relevant cardiac models are very limited. Humanized mouse models, while invaluable, cannot fully recapitulate the structural organization or electrophysiology properties of a human heart. Rodents exhibit different heart morphology, an approximatively tenfold higher heart rate and have an eightfold higher metabolic rate compared with humans ^76,77^. In this context, hIPSC-derived cardiomyocytes represent a critical model for cardiotoxicity assessment and drug efficacy testing ^40,43^. Although the incomplete maturation of hIPSC-derived cardiomyocytes has been documented ^80,81^, their spontaneous beating varies between batches (43 ±19 bpm). However, this range remains within the physiological range of human heart rate. As shown in this present study, spontaneous beating capability provides a physiologically relevant model to measure arrythmias and cardiac dysfunctions. Also, hIPSC-CMs are responsive to cellular injuries or pathogen exposure, triggering the secretion of inflammatory and cardioprotective markers including MIF, GRP78, ALDOA, STIP1, PDIA3, CFI, PGE 2 and PKM, as well as the release of heart failure-associated marker ^49–52,57,59^. In conclusion, the use of hIPSC-CMs is both relevant and reliable.

Manual counting and computational analysis were directionally consistent, but they captured distinct aspects of beating behavior. Manual counting provided an average beat rate over the recording interval and is well suited to detecting gross slowing or complete arrest. In contrast, the computational framework preserved beat-to-beat timing information and frequency content, enabling detection of irregular intervals, fragmented rhythmic components, and reduced motion amplitude that may precede complete beating arrest. Slight differences between the two approaches are therefore expected, particularly in traces with low-amplitude or irregular contractions, where manual counting may miss weak or closely spaced events and the algorithm may detect residual motion that is not readily apparent by visual inspection.

Several limitations of the computational approach should be acknowledged. First, the MAD signal is a full-field motion proxy and therefore reflects relative contractile activity rather than absolute contractile force, physical displacement, or electrophysiological state. Second, because the analysis is performed at the field-of-view level, local spatial heterogeneity or asynchronous subregions may contribute to the aggregate MAD trace. Third, the present validation was performed against manual beat counting, which supports beat detection accuracy but does not directly validate mechanical force generation or electrical conduction. Even with these limitations, the method provides a simple, non-invasive, and scalable approach for quantifying infection-induced functional decline from standard brightfield recordings.

## Author contributions

S.V.T., L.W., T.D., C.A., M.K. L.O., S.D.W., M.F.G, W.Z, Q.W., and E.R. performed experiments and analyzed data; S.V.T., L.M and T.D., A.N and Q.W., F.A. reviewed the manuscript; A.N., Q.W and E.R. conceived the study; E.R. led the study and wrote the manuscript.

## Supplementary Figures

**Supplementary Figure 1:**
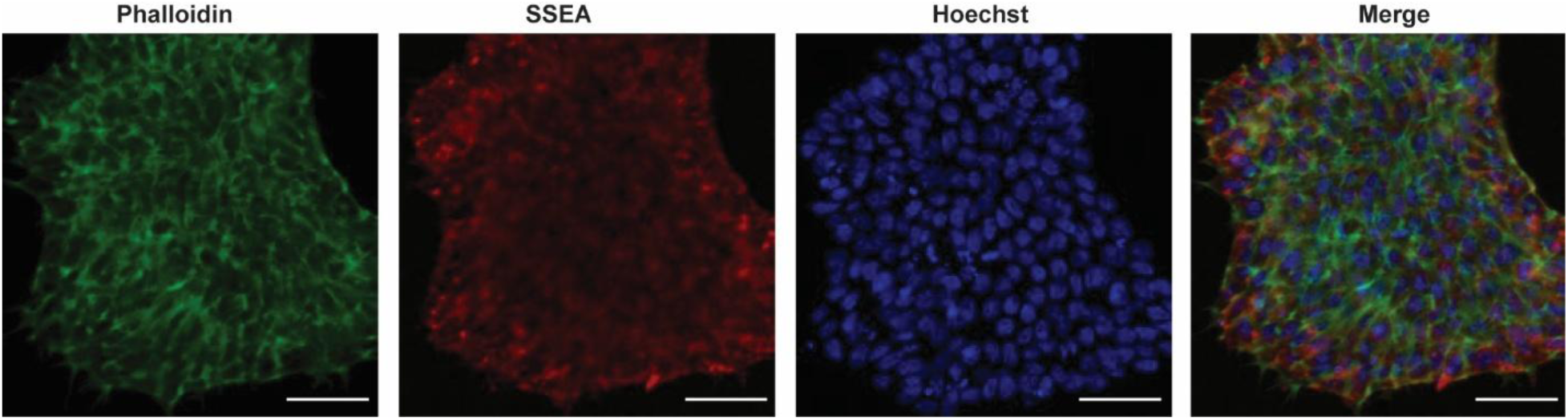
hIPSC characterization. Representative confocal images of Phalloidin, SSEA and Hoechst staining of hIPSC, scale bars = 50µm.

**Supplementary figure 2:**
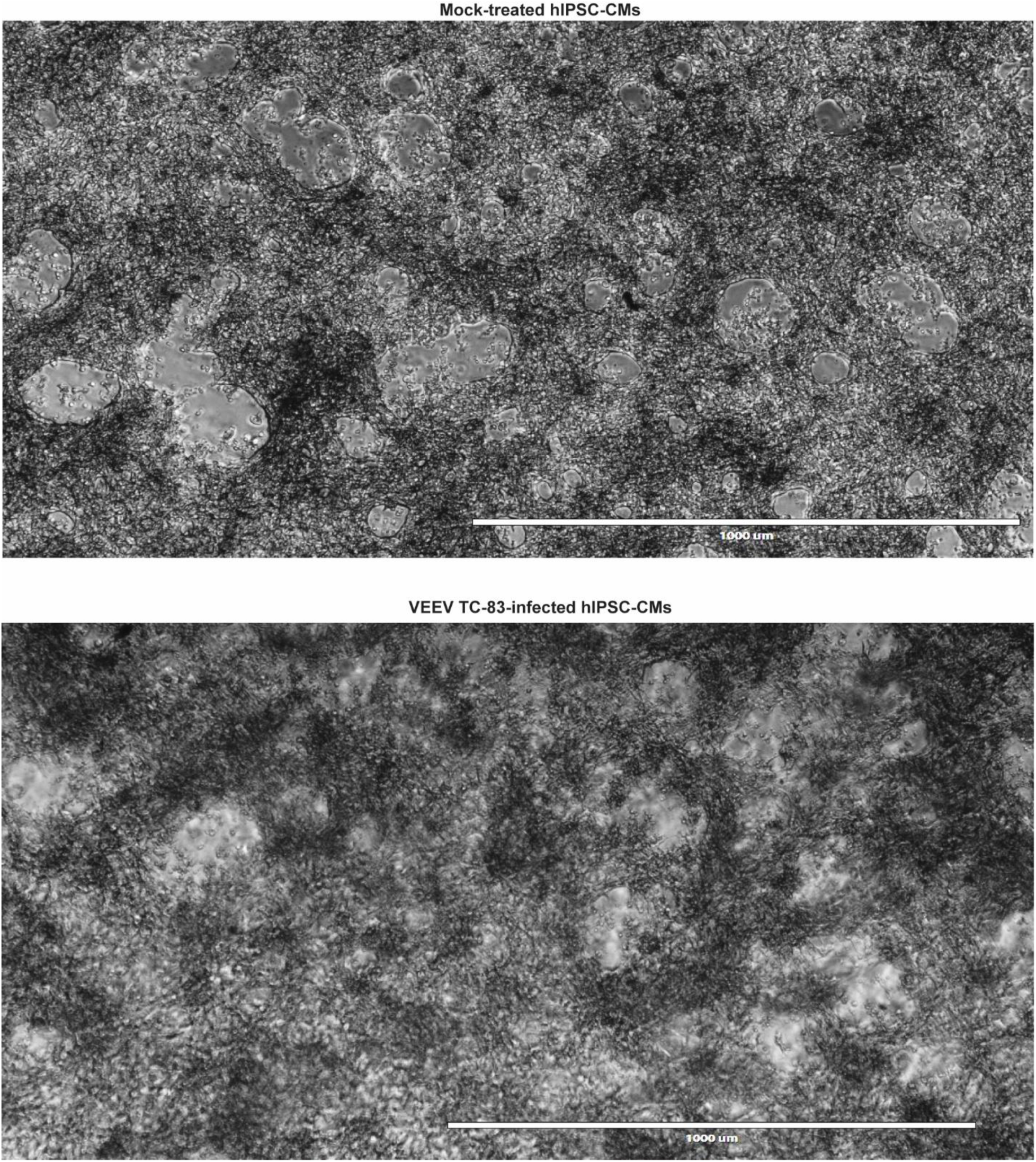
Representative brightfield images of hIPSC-derived cardiomyocytes mock-treated (top) or VEEV TC-83-infected (bottom) hIPSC-CMs at MOI 0.1, at 30 hpi. Scale bars = 1000 µm.

**Supplementary Figure 3:**
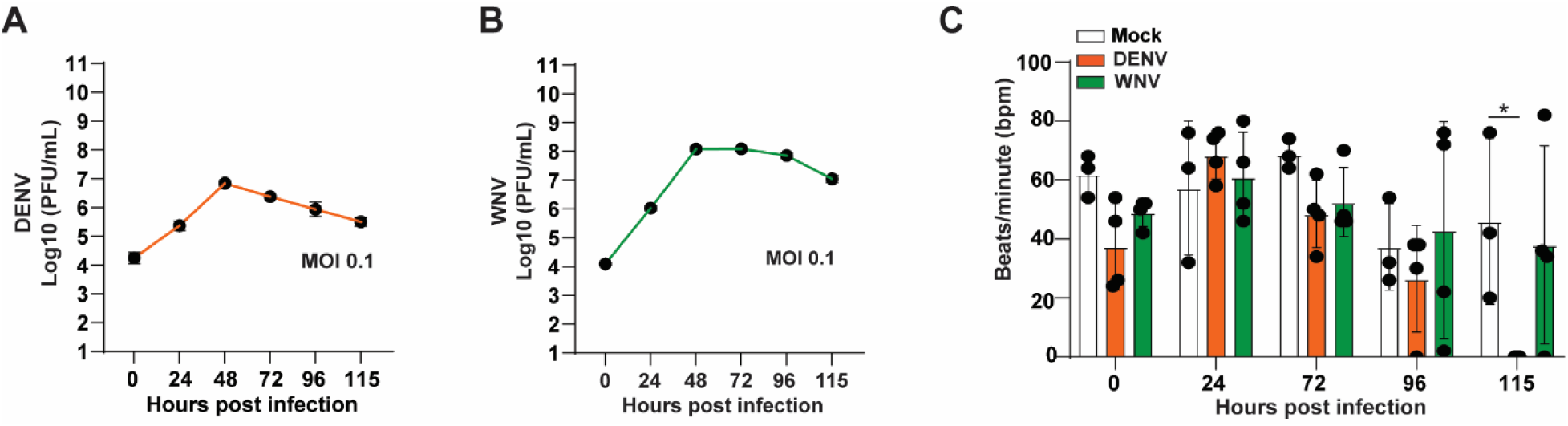
Assessment of Dengue and West Nile Virus susceptibility in Human Induced Pluripotent Stem Cell-derived Cardiomyocytes. A. Average of Dengue (DENV) titration (PFU/ml) upon hIPSC-CM infection. Infection at MOI 0.1 upon 115 hours, n=4 independently treated wells. B. Average of West Nile virus (WNV) titration (PFU/ml) upon hIPSC-CM infection, n=4 independently treated wells. C. Average of beating rhythm (beats/minute) of hIPSC-CM upon mock, DENV or WNV infection after T0, T24, T72, T96, and T115 hpi, n=4 independently treated wells, * p≤0.05.

**Supplementary Figure 4:**
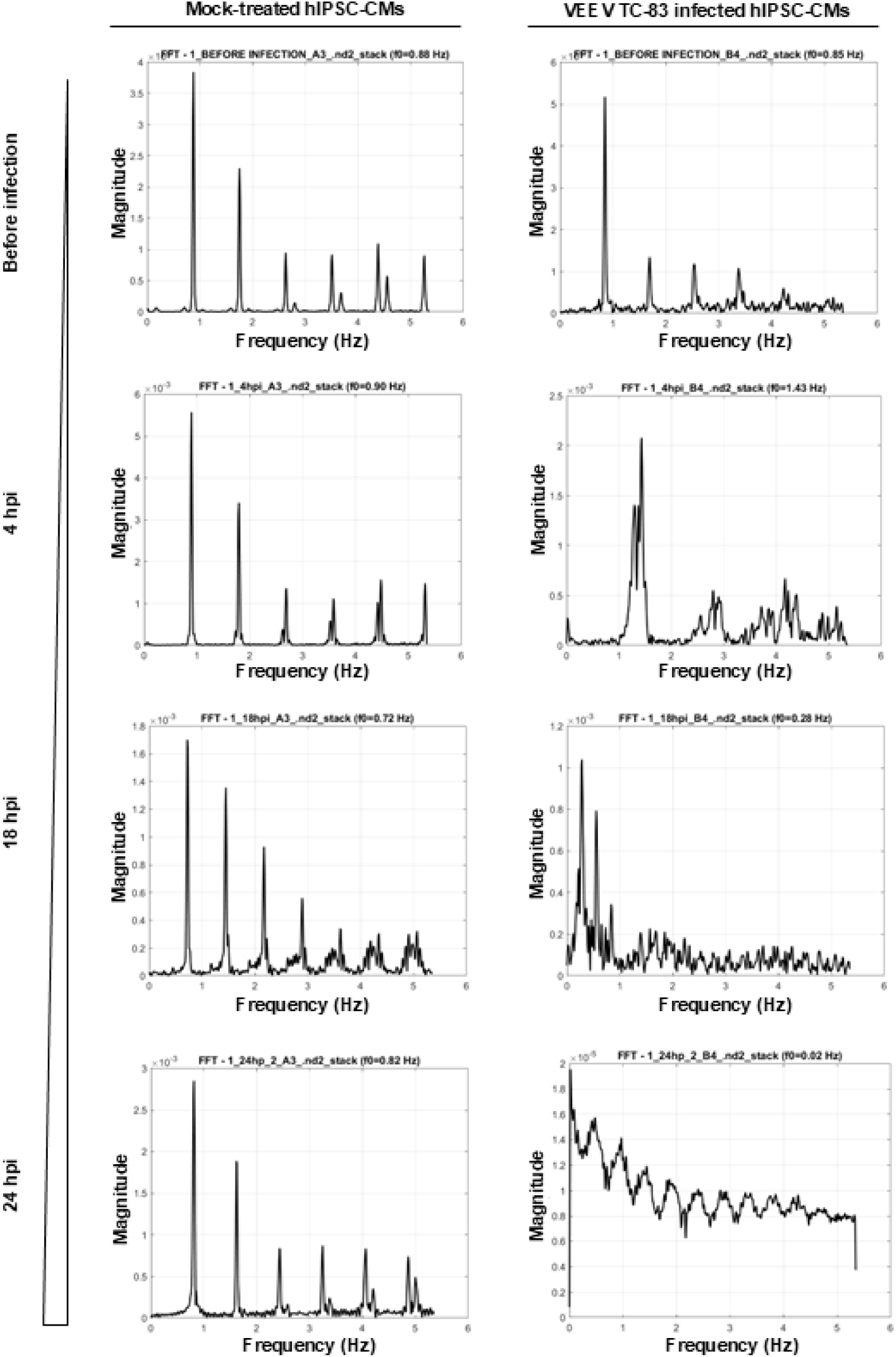
Frequency-domain analysis of MAD traces from mock-treated and VEEV TC-83-infected hIPSC-CMs. Representative fast Fourier transform (FFT) magnitude spectra and corresponding MAD traces are shown for mock-treated and infected cultures at T0, T4, T18, and T24 hpi. Mock-treated cultures retain a discrete dominant frequency across time points, whereas infected cultures show broadened, fragmented, or diminished spectral peaks consistent with rhythm instability and beating arrest.

**Supplementary Figure 5:**
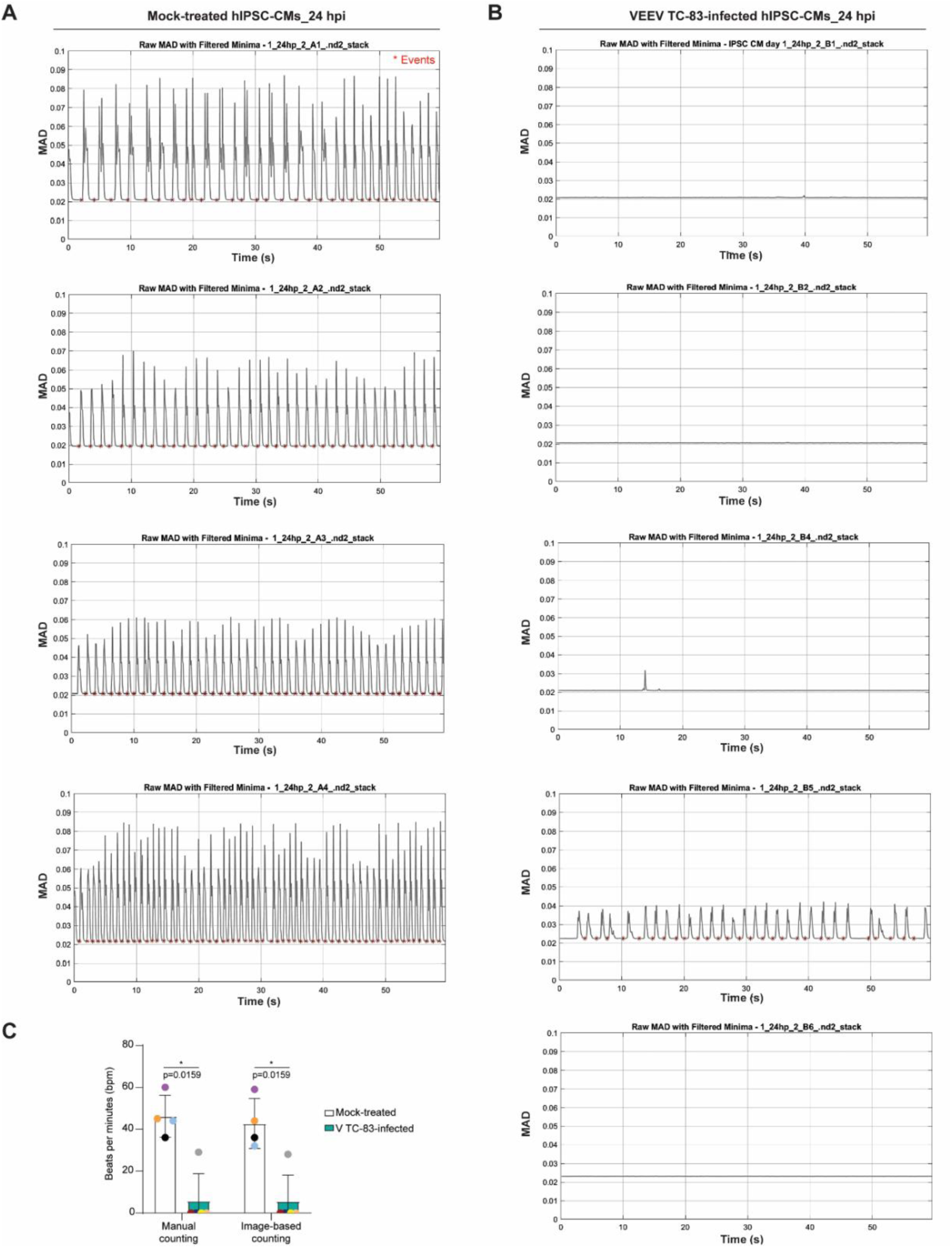
Reproducibility and comparison of manual versus computational beat analysis. A. Representative traces from independent mock-treated wells. B. Representative traces from independent VEEV TC-83-infected wells. C. Comparison between manually determined beat rate and algorithm-derived beat rate across wells and time points, showing strong agreement between the two methods.

**Supplementary Table 1:**
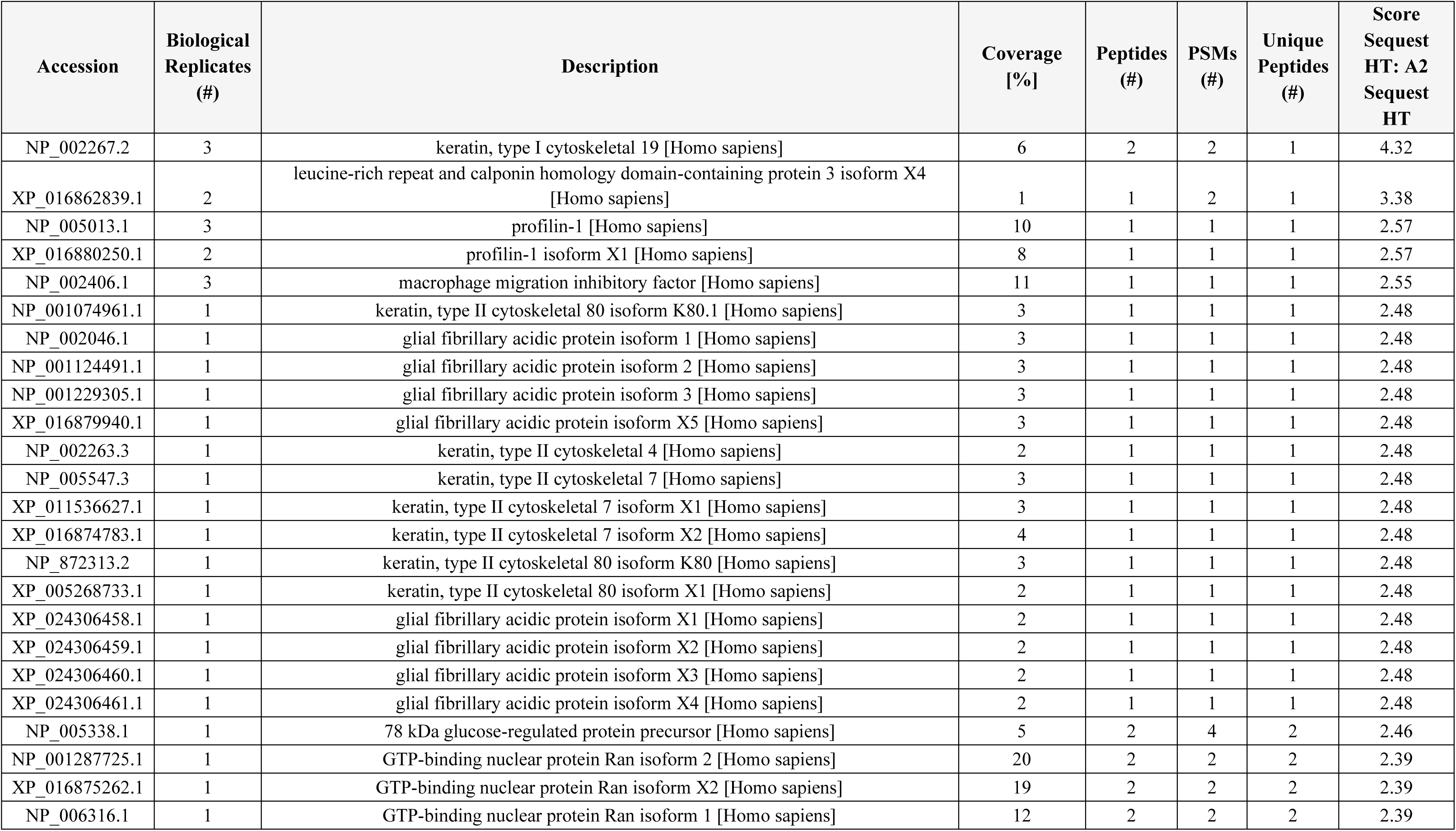
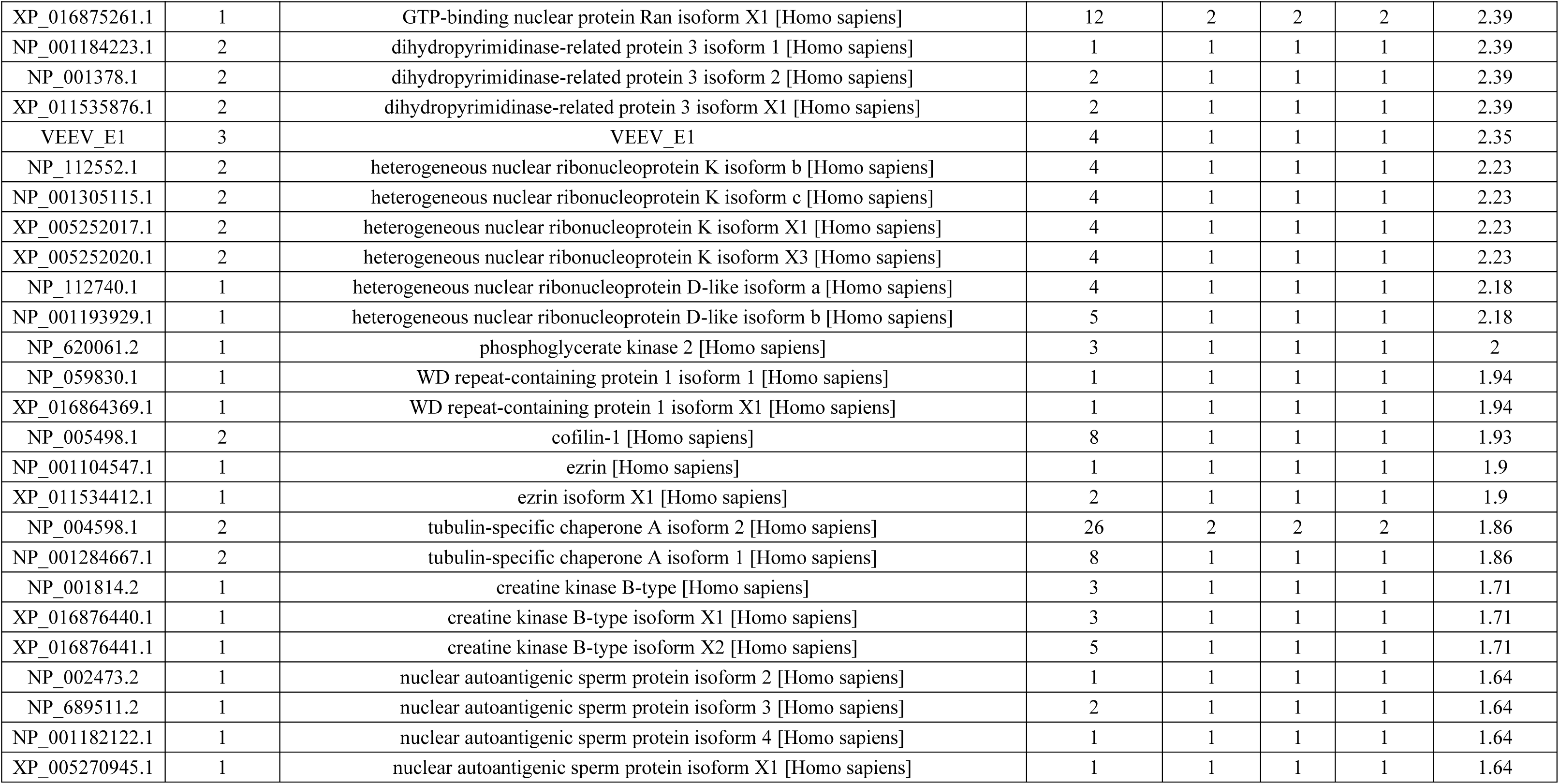
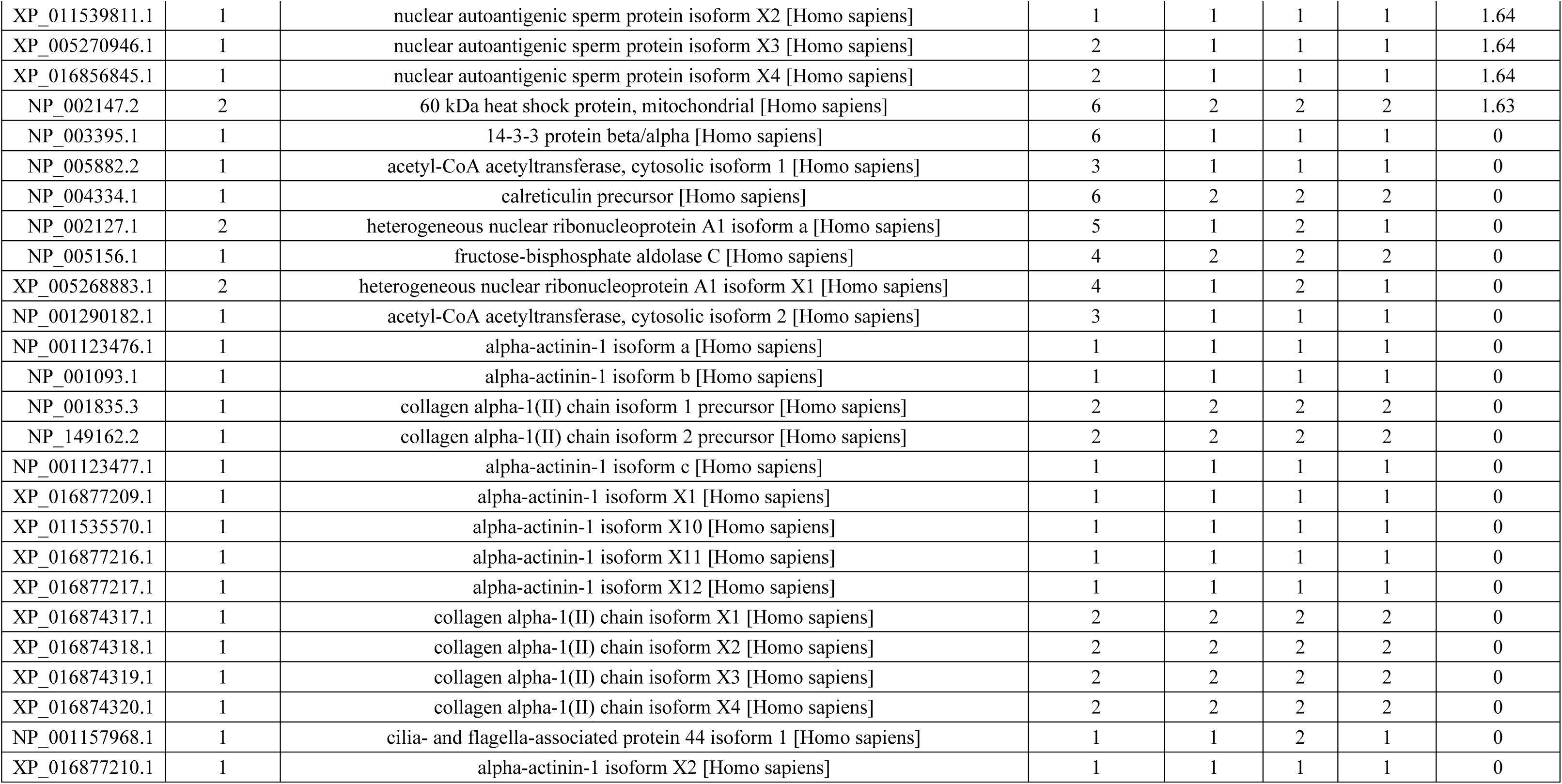
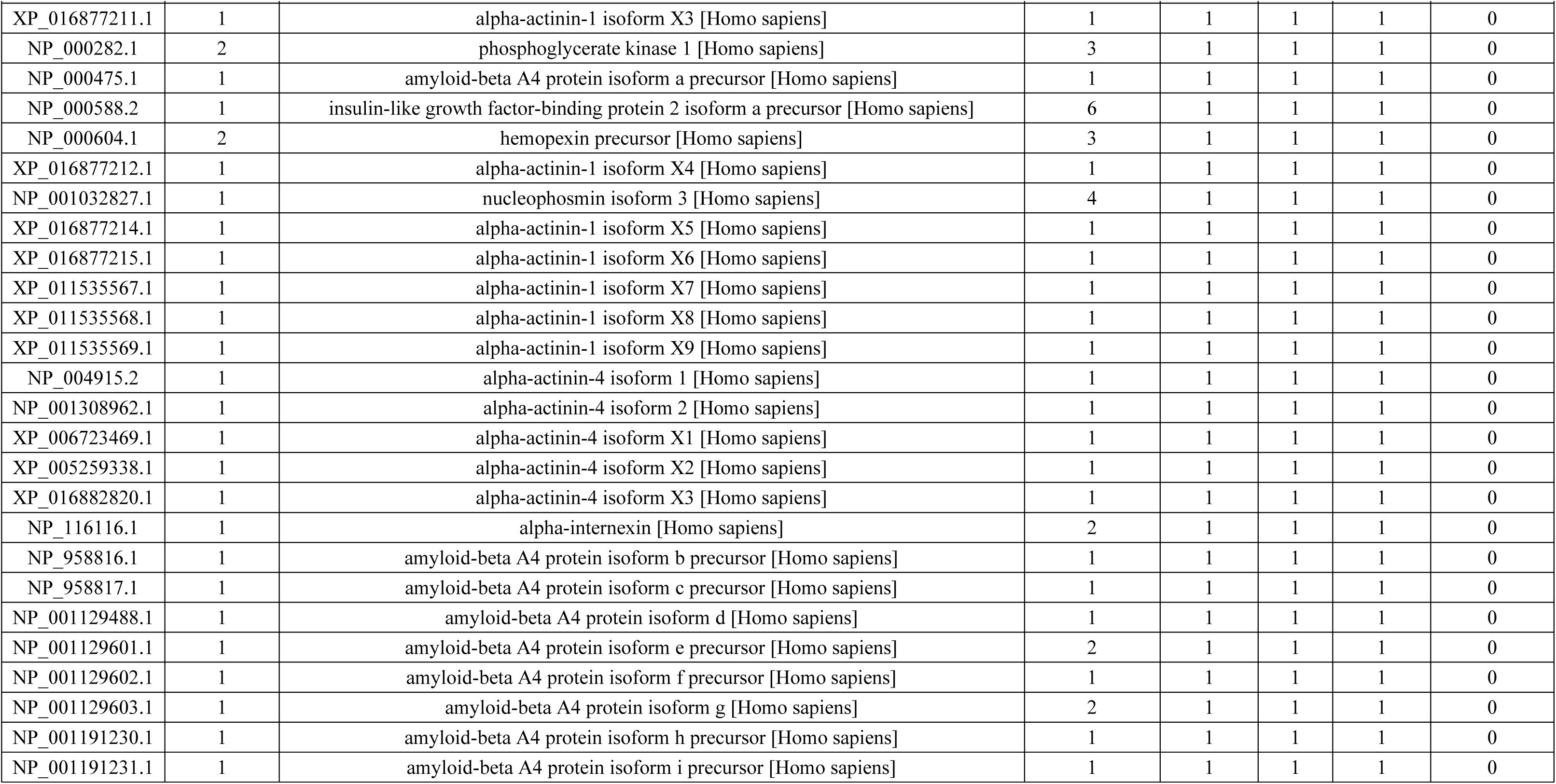
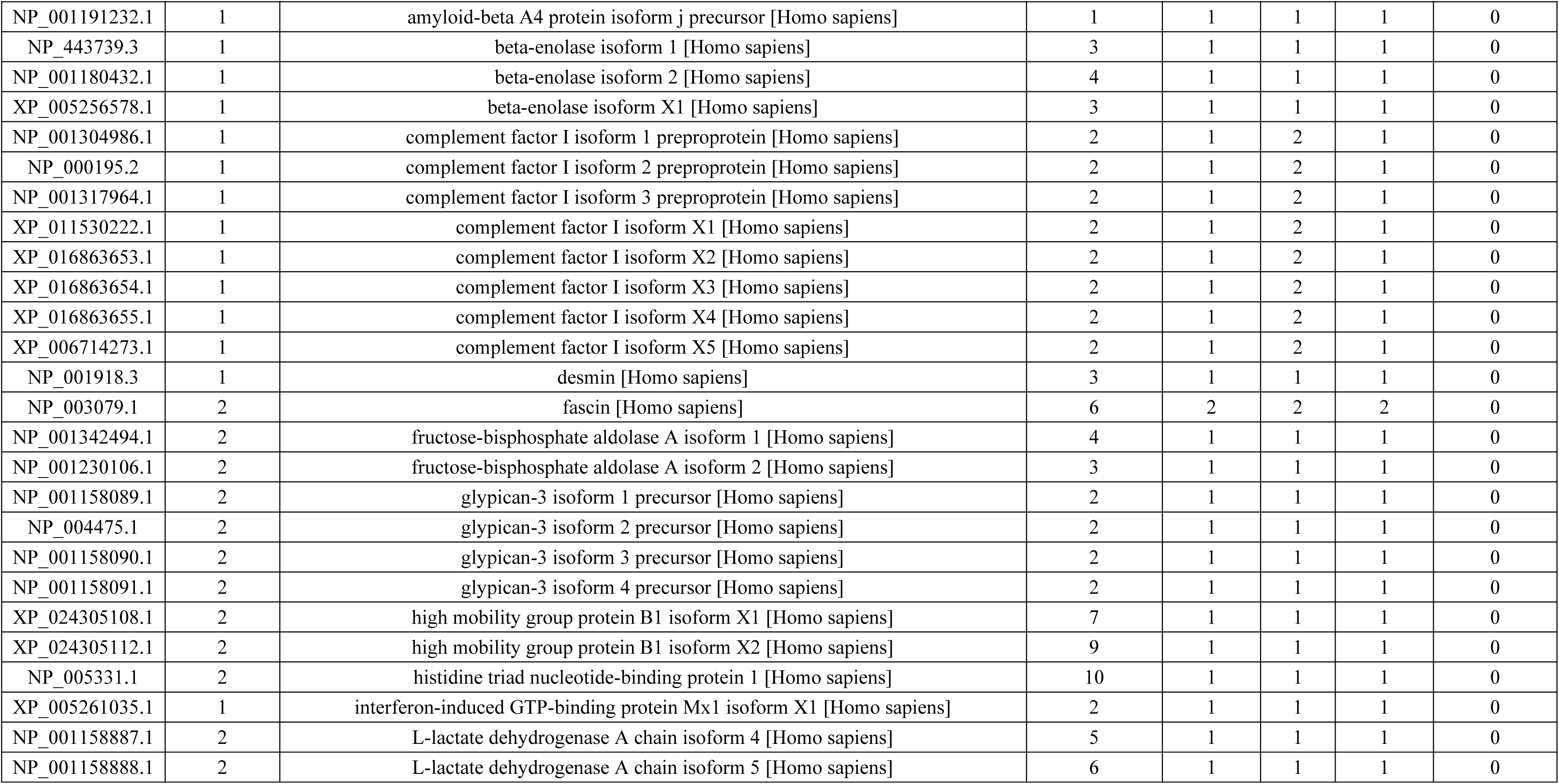
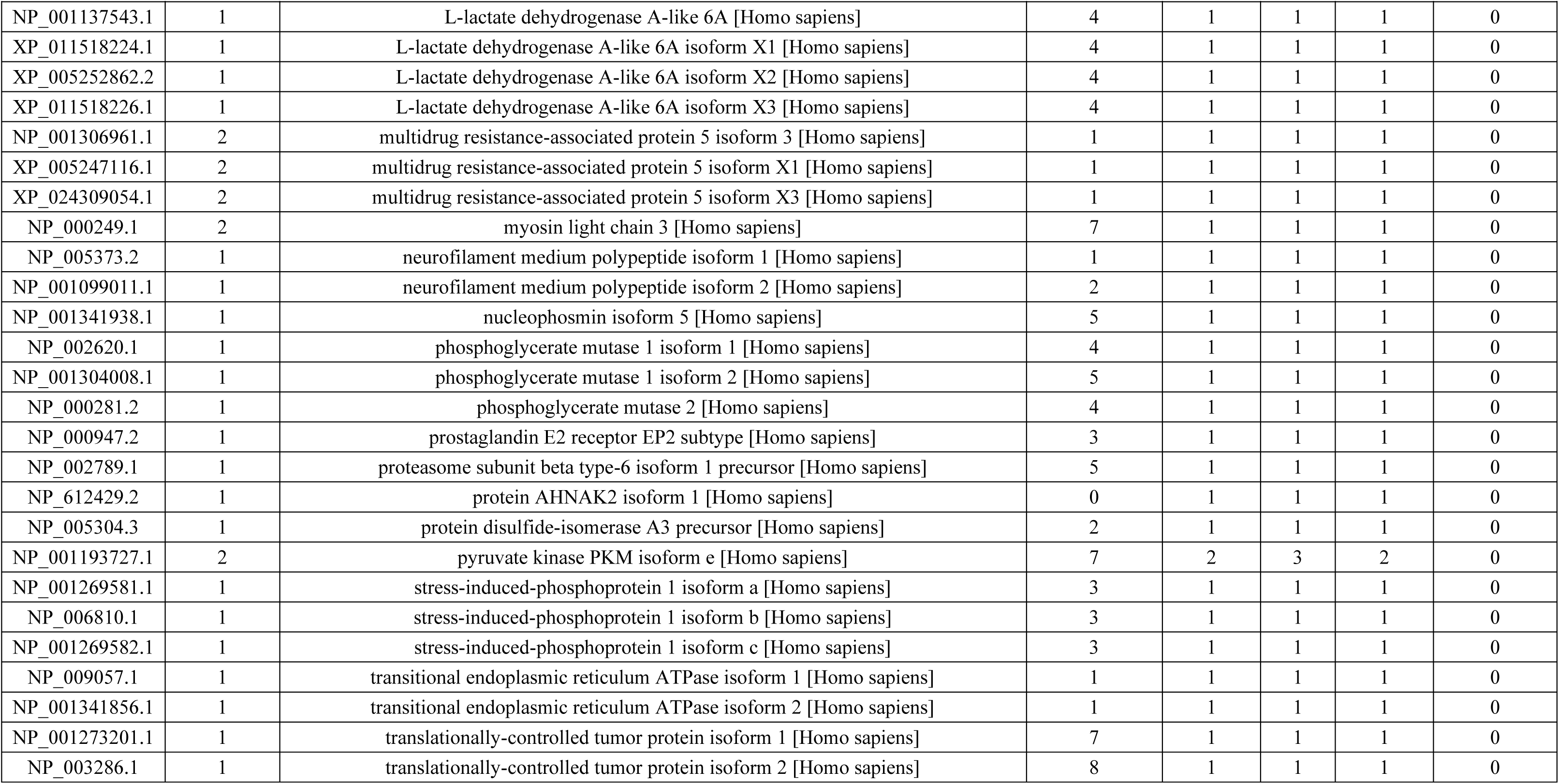
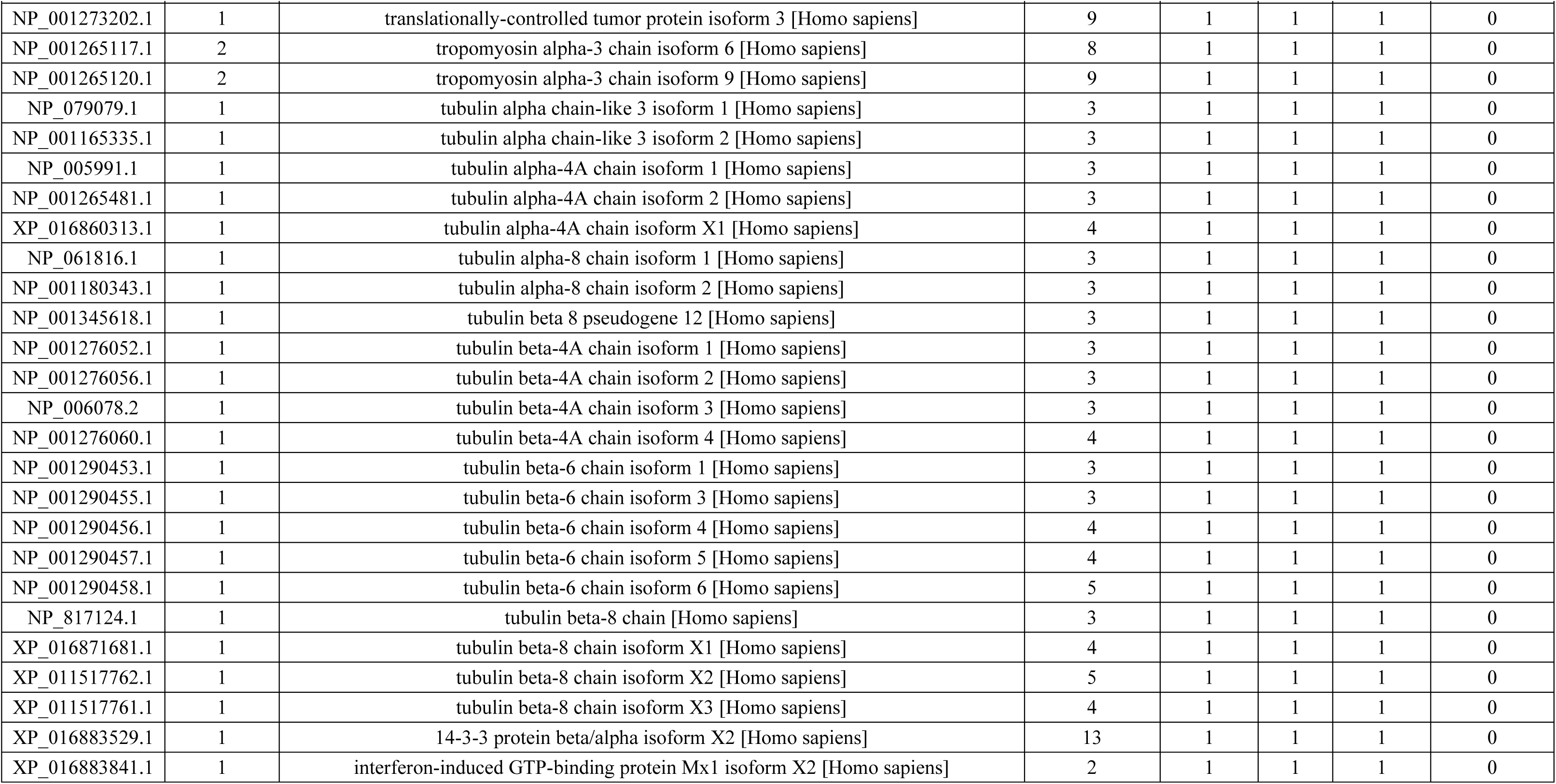
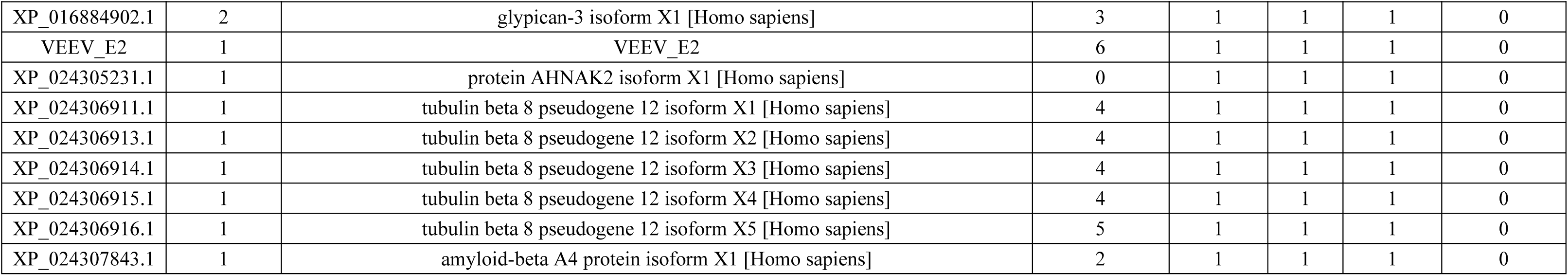
hIPSC-CM secreted Proteins Identified by LC-MS/MS specific to VEEV TC-83 infection. Table summarizes proteins identified by Liquid Chromatography-Tandem Mass Spectrometry (LC-MS/MS) from supernatant of VEEV TC-83 infected hIPSC-CMs (n = 3). Sequest score above medium confidence threshold, but below high confidence threshold reported as zero. PSM = peptide spectral matches.

| Accession | Biological Replicates (#) | Description | Coverage [%] | Peptides (#) | PSMs (#) | Unique Peptides (#) | Score<br>Sequest<br>HT: A2<br>Sequest<br>HT |
| --- | --- | --- | --- | --- | --- | --- | --- |
| NP_002267.2 | 3 | keratin, type I cytoskeletal 19 [Homo sapiens] | 6 | 2 | 2 | 1 | 4.32 |
| XP_016862839.1 | 2 | leucine-rich repeat and calponin homology domain-containing protein 3 isoform X4 [Homo sapiens] | 1 | 1 | 2 | 1 | 3.38 |
| NP_005013.1 | 3 | profilin-1 [Homo sapiens] | 10 | 1 | 1 | 1 | 2.57 |
| XP_016880250.1 | 2 | profilin-1 isoform X1 [Homo sapiens] | 8 | 1 | 1 | 1 | 2.57 |
| NP_002406.1 | 3 | macrophage migration inhibitory factor [Homo sapiens] | 11 | 1 | 1 | 1 | 2.55 |
| NP_001074961.1 | 1 | keratin, type II cytoskeletal 80 isoform K80.1 [Homo sapiens] | 3 | 1 | 1 | 1 | 2.48 |
| NP_002046.1 | 1 | glial fibrillary acidic protein isoform 1 [Homo sapiens] | 3 | 1 | 1 | 1 | 2.48 |
| NP_001124491.1 | 1 | glial fibrillary acidic protein isoform 2 [Homo sapiens] | 3 | 1 | 1 | 1 | 2.48 |
| NP_001229305.1 | 1 | glial fibrillary acidic protein isoform 3 [Homo sapiens] | 3 | 1 | 1 | 1 | 2.48 |
| XP_016879940.1 | 1 | glial fibrillary acidic protein isoform X5 [Homo sapiens] | 3 | 1 | 1 | 1 | 2.48 |
| NP_002263.3 | 1 | keratin, type II cytoskeletal 4 [Homo sapiens] | 2 | 1 | 1 | 1 | 2.48 |
| NP_005547.3 | 1 | keratin, type II cytoskeletal 7 [Homo sapiens] | 3 | 1 | 1 | 1 | 2.48 |
| XP_011536627.1 | 1 | keratin, type II cytoskeletal 7 isoform X1 [Homo sapiens] | 3 | 1 | 1 | 1 | 2.48 |
| XP_016874783.1 | 1 | keratin, type II cytoskeletal 7 isoform X2 [Homo sapiens] | 4 | 1 | 1 | 1 | 2.48 |
| NP_872313.2 | 1 | keratin, type II cytoskeletal 80 isoform K80 [Homo sapiens] | 3 | 1 | 1 | 1 | 2.48 |
| XP_005268733.1 | 1 | keratin, type II cytoskeletal 80 isoform X1 [Homo sapiens] | 2 | 1 | 1 | 1 | 2.48 |
| XP_024306458.1 | 1 | glial fibrillary acidic protein isoform X1 [Homo sapiens] | 2 | 1 | 1 | 1 | 2.48 |
| XP_024306459.1 | 1 | glial fibrillary acidic protein isoform X2 [Homo sapiens] | 2 | 1 | 1 | 1 | 2.48 |
| XP_024306460.1 | 1 | glial fibrillary acidic protein isoform X3 [Homo sapiens] | 2 | 1 | 1 | 1 | 2.48 |
| XP_024306461.1 | 1 | glial fibrillary acidic protein isoform X4 [Homo sapiens] | 2 | 1 | 1 | 1 | 2.48 |
| NP_005338.1 | 1 | 78 kDa glucose-regulated protein precursor [Homo sapiens] | 5 | 2 | 4 | 2 | 2.46 |
| NP_001287725.1 | 1 | GTP-binding nuclear protein Ran isoform 2 [Homo sapiens] | 20 | 2 | 2 | 2 | 2.39 |
| XP_016875262.1 | 1 | GTP-binding nuclear protein Ran isoform X2 [Homo sapiens] | 19 | 2 | 2 | 2 | 2.39 |
| NP_006316.1 | 1 | GTP-binding nuclear protein Ran isoform 1 [Homo sapiens] | 12 | 2 | 2 | 2 | 2.39 |

| Accession | Biological Replicates (#) | Description | Coverage [%] | Peptides (#) | PSMs (#) | Unique Peptides (#) | Score<br>Sequest HT: A2<br>Sequest HT |
| --- | --- | --- | --- | --- | --- | --- | --- |
| XP_016875261.1 | 1 | GTP-binding nuclear protein Ran isoform X1 [Homo sapiens] | 12 | 2 | 2 | 2 | 2.39 |
| NP_001184223.1 | 2 | dihydropyrimidinase-related protein 3 isoform 1 [Homo sapiens] | 1 | 1 | 1 | 1 | 2.39 |
| NP_001378.1 | 2 | dihydropyrimidinase-related protein 3 isoform 2 [Homo sapiens] | 2 | 1 | 1 | 1 | 2.39 |
| XP_011535876.1 | 2 | dihydropyrimidinase-related protein 3 isoform X1 [Homo sapiens] | 2 | 1 | 1 | 1 | 2.39 |
| VEEV_E1 | 3 | VEEV_E1 | 4 | 1 | 1 | 1 | 2.35 |
| NP_112552.1 | 2 | heterogeneous nuclear ribonucleoprotein K isoform b [Homo sapiens] | 4 | 1 | 1 | 1 | 2.23 |
| NP_001305115.1 | 2 | heterogeneous nuclear ribonucleoprotein K isoform c [Homo sapiens] | 4 | 1 | 1 | 1 | 2.23 |
| XP_005252017.1 | 2 | heterogeneous nuclear ribonucleoprotein K isoform X1 [Homo sapiens] | 4 | 1 | 1 | 1 | 2.23 |
| XP_005252020.1 | 2 | heterogeneous nuclear ribonucleoprotein K isoform X3 [Homo sapiens] | 4 | 1 | 1 | 1 | 2.23 |
| NP_112740.1 | 1 | heterogeneous nuclear ribonucleoprotein D-like isoform a [Homo sapiens] | 4 | 1 | 1 | 1 | 2.18 |
| NP_001193929.1 | 1 | heterogeneous nuclear ribonucleoprotein D-like isoform b [Homo sapiens] | 5 | 1 | 1 | 1 | 2.18 |
| NP_620061.2 | 1 | phosphoglycerate kinase 2 [Homo sapiens] | 3 | 1 | 1 | 1 | 2 |
| NP_059830.1 | 1 | WD repeat-containing protein 1 isoform 1 [Homo sapiens] | 1 | 1 | 1 | 1 | 1.94 |
| XP_016864369.1 | 1 | WD repeat-containing protein 1 isoform X1 [Homo sapiens] | 1 | 1 | 1 | 1 | 1.94 |
| NP_005498.1 | 2 | cofilin-1 [Homo sapiens] | 8 | 1 | 1 | 1 | 1.93 |
| NP_001104547.1 | 1 | ezrin [Homo sapiens] | 1 | 1 | 1 | 1 | 1.9 |
| XP_011534412.1 | 1 | ezrin isoform X1 [Homo sapiens] | 2 | 1 | 1 | 1 | 1.9 |
| NP_004598.1 | 2 | tubulin-specific chaperone A isoform 2 [Homo sapiens] | 26 | 2 | 2 | 2 | 1.86 |
| NP_001284667.1 | 2 | tubulin-specific chaperone A isoform 1 [Homo sapiens] | 8 | 1 | 1 | 1 | 1.86 |
| NP_001814.2 | 1 | creatine kinase B-type [Homo sapiens] | 3 | 1 | 1 | 1 | 1.71 |
| XP_016876440.1 | 1 | creatine kinase B-type isoform X1 [Homo sapiens] | 3 | 1 | 1 | 1 | 1.71 |
| XP_016876441.1 | 1 | creatine kinase B-type isoform X2 [Homo sapiens] | 5 | 1 | 1 | 1 | 1.71 |
| NP_002473.2 | 1 | nuclear autoantigenic sperm protein isoform 2 [Homo sapiens] | 1 | 1 | 1 | 1 | 1.64 |
| NP_689511.2 | 1 | nuclear autoantigenic sperm protein isoform 3 [Homo sapiens] | 2 | 1 | 1 | 1 | 1.64 |
| NP_001182122.1 | 1 | nuclear autoantigenic sperm protein isoform 4 [Homo sapiens] | 1 | 1 | 1 | 1 | 1.64 |
| XP_005270945.1 | 1 | nuclear autoantigenic sperm protein isoform X1 [Homo sapiens] | 1 | 1 | 1 | 1 | 1.64 |

| Accession | Biological Replicates (#) | Description | Coverage [%] | Peptides (#) | PSMs (#) | Unique Peptides (#) | Score Sequest HT: A2 Sequest HT |
| --- | --- | --- | --- | --- | --- | --- | --- |
| XP_011539811.1 | 1 | nuclear autoantigenic sperm protein isoform X2 [Homo sapiens] | 1 | 1 | 1 | 1 | 1.64 |
| XP_005270946.1 | 1 | nuclear autoantigenic sperm protein isoform X3 [Homo sapiens] | 2 | 1 | 1 | 1 | 1.64 |
| XP_016856845.1 | 1 | nuclear autoantigenic sperm protein isoform X4 [Homo sapiens] | 2 | 1 | 1 | 1 | 1.64 |
| NP_002147.2 | 2 | 60 kDa heat shock protein, mitochondrial [Homo sapiens] | 6 | 2 | 2 | 2 | 1.63 |
| NP_003395.1 | 1 | 14-3-3 protein beta/alpha [Homo sapiens] | 6 | 1 | 1 | 1 | 0 |
| NP_005882.2 | 1 | acetyl-CoA acetyltransferase, cytosolic isoform 1 [Homo sapiens] | 3 | 1 | 1 | 1 | 0 |
| NP_004334.1 | 1 | calreticulin precursor [Homo sapiens] | 6 | 2 | 2 | 2 | 0 |
| NP_002127.1 | 2 | heterogeneous nuclear ribonucleoprotein A1 isoform a [Homo sapiens] | 5 | 1 | 2 | 1 | 0 |
| NP_005156.1 | 1 | fructose-bisphosphate aldolase C [Homo sapiens] | 4 | 2 | 2 | 2 | 0 |
| XP_005268883.1 | 2 | heterogeneous nuclear ribonucleoprotein A1 isoform X1 [Homo sapiens] | 4 | 1 | 2 | 1 | 0 |
| NP_001290182.1 | 1 | acetyl-CoA acetyltransferase, cytosolic isoform 2 [Homo sapiens] | 3 | 1 | 1 | 1 | 0 |
| NP_001123476.1 | 1 | alpha-actinin-1 isoform a [Homo sapiens] | 1 | 1 | 1 | 1 | 0 |
| NP_001093.1 | 1 | alpha-actinin-1 isoform b [Homo sapiens] | 1 | 1 | 1 | 1 | 0 |
| NP_001835.3 | 1 | collagen alpha-1(II) chain isoform 1 precursor [Homo sapiens] | 2 | 2 | 2 | 2 | 0 |
| NP_149162.2 | 1 | collagen alpha-1(II) chain isoform 2 precursor [Homo sapiens] | 2 | 2 | 2 | 2 | 0 |
| NP_001123477.1 | 1 | alpha-actinin-1 isoform c [Homo sapiens] | 1 | 1 | 1 | 1 | 0 |
| XP_016877209.1 | 1 | alpha-actinin-1 isoform X1 [Homo sapiens] | 1 | 1 | 1 | 1 | 0 |
| XP_011535570.1 | 1 | alpha-actinin-1 isoform X10 [Homo sapiens] | 1 | 1 | 1 | 1 | 0 |
| XP_016877216.1 | 1 | alpha-actinin-1 isoform X11 [Homo sapiens] | 1 | 1 | 1 | 1 | 0 |
| XP_016877217.1 | 1 | alpha-actinin-1 isoform X12 [Homo sapiens] | 1 | 1 | 1 | 1 | 0 |
| XP_016874317.1 | 1 | collagen alpha-1(II) chain isoform X1 [Homo sapiens] | 2 | 2 | 2 | 2 | 0 |
| XP_016874318.1 | 1 | collagen alpha-1(II) chain isoform X2 [Homo sapiens] | 2 | 2 | 2 | 2 | 0 |
| XP_016874319.1 | 1 | collagen alpha-1(II) chain isoform X3 [Homo sapiens] | 2 | 2 | 2 | 2 | 0 |
| XP_016874320.1 | 1 | collagen alpha-1(II) chain isoform X4 [Homo sapiens] | 2 | 2 | 2 | 2 | 0 |
| NP_001157968.1 | 1 | cilia- and flagella-associated protein 44 isoform 1 [Homo sapiens] | 1 | 1 | 2 | 1 | 0 |
| XP_016877210.1 | 1 | alpha-actinin-1 isoform X2 [Homo sapiens] | 1 | 1 | 1 | 1 | 0 |
| XP_016877211.1 | 1 | alpha-actinin-1 isoform X3 [Homo sapiens] | 1 | 1 | 1 | 1 | 0 |
| NP_000282.1 | 2 | phosphoglycerate kinase 1 [Homo sapiens] | 3 | 1 | 1 | 1 | 0 |
| NP_000475.1 | 1 | amyloid-beta A4 protein isoform a precursor [Homo sapiens] | 1 | 1 | 1 | 1 | 0 |
| NP_000588.2 | 1 | insulin-like growth factor-binding protein 2 isoform a precursor [Homo sapiens] | 6 | 1 | 1 | 1 | 0 |
| NP_000604.1 | 2 | hemopexin precursor [Homo sapiens] | 3 | 1 | 1 | 1 | 0 |
| XP_016877212.1 | 1 | alpha-actinin-1 isoform X4 [Homo sapiens] | 1 | 1 | 1 | 1 | 0 |
| NP_001032827.1 | 1 | nucleophosmin isoform 3 [Homo sapiens] | 4 | 1 | 1 | 1 | 0 |
| XP_016877214.1 | 1 | alpha-actinin-1 isoform X5 [Homo sapiens] | 1 | 1 | 1 | 1 | 0 |
| XP_016877215.1 | 1 | alpha-actinin-1 isoform X6 [Homo sapiens] | 1 | 1 | 1 | 1 | 0 |
| XP_011535567.1 | 1 | alpha-actinin-1 isoform X7 [Homo sapiens] | 1 | 1 | 1 | 1 | 0 |
| XP_011535568.1 | 1 | alpha-actinin-1 isoform X8 [Homo sapiens] | 1 | 1 | 1 | 1 | 0 |
| XP_011535569.1 | 1 | alpha-actinin-1 isoform X9 [Homo sapiens] | 1 | 1 | 1 | 1 | 0 |
| NP_004915.2 | 1 | alpha-actinin-4 isoform 1 [Homo sapiens] | 1 | 1 | 1 | 1 | 0 |
| NP_001308962.1 | 1 | alpha-actinin-4 isoform 2 [Homo sapiens] | 1 | 1 | 1 | 1 | 0 |
| XP_006723469.1 | 1 | alpha-actinin-4 isoform X1 [Homo sapiens] | 1 | 1 | 1 | 1 | 0 |
| XP_005259338.1 | 1 | alpha-actinin-4 isoform X2 [Homo sapiens] | 1 | 1 | 1 | 1 | 0 |
| XP_016882820.1 | 1 | alpha-actinin-4 isoform X3 [Homo sapiens] | 1 | 1 | 1 | 1 | 0 |
| NP_116116.1 | 1 | alpha-internexin [Homo sapiens] | 2 | 1 | 1 | 1 | 0 |
| NP_958816.1 | 1 | amyloid-beta A4 protein isoform b precursor [Homo sapiens] | 1 | 1 | 1 | 1 | 0 |
| NP_958817.1 | 1 | amyloid-beta A4 protein isoform c precursor [Homo sapiens] | 1 | 1 | 1 | 1 | 0 |
| NP_001129488.1 | 1 | amyloid-beta A4 protein isoform d [Homo sapiens] | 1 | 1 | 1 | 1 | 0 |
| NP_001129601.1 | 1 | amyloid-beta A4 protein isoform e precursor [Homo sapiens] | 2 | 1 | 1 | 1 | 0 |
| NP_001129602.1 | 1 | amyloid-beta A4 protein isoform f precursor [Homo sapiens] | 1 | 1 | 1 | 1 | 0 |
| NP_001129603.1 | 1 | amyloid-beta A4 protein isoform g [Homo sapiens] | 2 | 1 | 1 | 1 | 0 |
| NP_001191230.1 | 1 | amyloid-beta A4 protein isoform h precursor [Homo sapiens] | 1 | 1 | 1 | 1 | 0 |
| NP_001191231.1 | 1 | amyloid-beta A4 protein isoform i precursor [Homo sapiens] | 1 | 1 | 1 | 1 | 0 |
| NP_001191232.1 | 1 | amyloid-beta A4 protein isoform j precursor [Homo sapiens] | 1 | 1 | 1 | 1 | 0 |
| NP_443739.3 | 1 | beta-enolase isoform 1 [Homo sapiens] | 3 | 1 | 1 | 1 | 0 |
| NP_001180432.1 | 1 | beta-enolase isoform 2 [Homo sapiens] | 4 | 1 | 1 | 1 | 0 |
| XP_005256578.1 | 1 | beta-enolase isoform X1 [Homo sapiens] | 3 | 1 | 1 | 1 | 0 |
| NP_001304986.1 | 1 | complement factor I isoform 1 preproprotein [Homo sapiens] | 2 | 1 | 2 | 1 | 0 |
| NP_000195.2 | 1 | complement factor I isoform 2 preproprotein [Homo sapiens] | 2 | 1 | 2 | 1 | 0 |
| NP_001317964.1 | 1 | complement factor I isoform 3 preproprotein [Homo sapiens] | 2 | 1 | 2 | 1 | 0 |
| XP_011530222.1 | 1 | complement factor I isoform X1 [Homo sapiens] | 2 | 1 | 2 | 1 | 0 |
| XP_016863653.1 | 1 | complement factor I isoform X2 [Homo sapiens] | 2 | 1 | 2 | 1 | 0 |
| XP_016863654.1 | 1 | complement factor I isoform X3 [Homo sapiens] | 2 | 1 | 2 | 1 | 0 |
| XP_016863655.1 | 1 | complement factor I isoform X4 [Homo sapiens] | 2 | 1 | 2 | 1 | 0 |
| XP_006714273.1 | 1 | complement factor I isoform X5 [Homo sapiens] | 2 | 1 | 2 | 1 | 0 |
| NP_001918.3 | 1 | desmin [Homo sapiens] | 3 | 1 | 1 | 1 | 0 |
| NP_003079.1 | 2 | fascin [Homo sapiens] | 6 | 2 | 2 | 2 | 0 |
| NP_001342494.1 | 2 | fructose-bisphosphate aldolase A isoform 1 [Homo sapiens] | 4 | 1 | 1 | 1 | 0 |
| NP_001230106.1 | 2 | fructose-bisphosphate aldolase A isoform 2 [Homo sapiens] | 3 | 1 | 1 | 1 | 0 |
| NP_001158089.1 | 2 | glypican-3 isoform 1 precursor [Homo sapiens] | 2 | 1 | 1 | 1 | 0 |
| NP_004475.1 | 2 | glypican-3 isoform 2 precursor [Homo sapiens] | 2 | 1 | 1 | 1 | 0 |
| NP_001158090.1 | 2 | glypican-3 isoform 3 precursor [Homo sapiens] | 2 | 1 | 1 | 1 | 0 |
| NP_001158091.1 | 2 | glypican-3 isoform 4 precursor [Homo sapiens] | 2 | 1 | 1 | 1 | 0 |
| XP_024305108.1 | 2 | high mobility group protein B1 isoform X1 [Homo sapiens] | 7 | 1 | 1 | 1 | 0 |
| XP_024305112.1 | 2 | high mobility group protein B1 isoform X2 [Homo sapiens] | 9 | 1 | 1 | 1 | 0 |
| NP_005331.1 | 2 | histidine triad nucleotide-binding protein 1 [Homo sapiens] | 10 | 1 | 1 | 1 | 0 |
| XP_005261035.1 | 1 | interferon-induced GTP-binding protein Mx1 isoform X1 [Homo sapiens] | 2 | 1 | 1 | 1 | 0 |
| NP_001158887.1 | 2 | L-lactate dehydrogenase A chain isoform 4 [Homo sapiens] | 5 | 1 | 1 | 1 | 0 |
| NP_001158888.1 | 2 | L-lactate dehydrogenase A chain isoform 5 [Homo sapiens] | 6 | 1 | 1 | 1 | 0 |
| NP_001137543.1 | 1 | L-lactate dehydrogenase A-like 6A [Homo sapiens] | 4 | 1 | 1 | 1 | 0 |
| XP_011518224.1 | 1 | L-lactate dehydrogenase A-like 6A isoform X1 [Homo sapiens] | 4 | 1 | 1 | 1 | 0 |
| XP_005252862.2 | 1 | L-lactate dehydrogenase A-like 6A isoform X2 [Homo sapiens] | 4 | 1 | 1 | 1 | 0 |
| XP_011518226.1 | 1 | L-lactate dehydrogenase A-like 6A isoform X3 [Homo sapiens] | 4 | 1 | 1 | 1 | 0 |
| NP_001306961.1 | 2 | multidrug resistance-associated protein 5 isoform 3 [Homo sapiens] | 1 | 1 | 1 | 1 | 0 |
| XP_005247116.1 | 2 | multidrug resistance-associated protein 5 isoform X1 [Homo sapiens] | 1 | 1 | 1 | 1 | 0 |
| XP_024309054.1 | 2 | multidrug resistance-associated protein 5 isoform X3 [Homo sapiens] | 1 | 1 | 1 | 1 | 0 |
| NP_000249.1 | 2 | myosin light chain 3 [Homo sapiens] | 7 | 1 | 1 | 1 | 0 |
| NP_005373.2 | 1 | neurofilament medium polypeptide isoform 1 [Homo sapiens] | 1 | 1 | 1 | 1 | 0 |
| NP_001099011.1 | 1 | neurofilament medium polypeptide isoform 2 [Homo sapiens] | 2 | 1 | 1 | 1 | 0 |
| NP_001341938.1 | 1 | nucleophosmin isoform 5 [Homo sapiens] | 5 | 1 | 1 | 1 | 0 |
| NP_002620.1 | 1 | phosphoglycerate mutase 1 isoform 1 [Homo sapiens] | 4 | 1 | 1 | 1 | 0 |
| NP_001304008.1 | 1 | phosphoglycerate mutase 1 isoform 2 [Homo sapiens] | 5 | 1 | 1 | 1 | 0 |
| NP_000281.2 | 1 | phosphoglycerate mutase 2 [Homo sapiens] | 4 | 1 | 1 | 1 | 0 |
| NP_000947.2 | 1 | prostaglandin E2 receptor EP2 subtype [Homo sapiens] | 3 | 1 | 1 | 1 | 0 |
| NP_002789.1 | 1 | proteasome subunit beta type-6 isoform 1 precursor [Homo sapiens] | 5 | 1 | 1 | 1 | 0 |
| NP_612429.2 | 1 | protein AHNAK2 isoform 1 [Homo sapiens] | 0 | 1 | 1 | 1 | 0 |
| NP_005304.3 | 1 | protein disulfide-isomerase A3 precursor [Homo sapiens] | 2 | 1 | 1 | 1 | 0 |
| NP_001193727.1 | 2 | pyruvate kinase PKM isoform e [Homo sapiens] | 7 | 2 | 3 | 2 | 0 |
| NP_001269581.1 | 1 | stress-induced-phosphoprotein 1 isoform a [Homo sapiens] | 3 | 1 | 1 | 1 | 0 |
| NP_006810.1 | 1 | stress-induced-phosphoprotein 1 isoform b [Homo sapiens] | 3 | 1 | 1 | 1 | 0 |
| NP_001269582.1 | 1 | stress-induced-phosphoprotein 1 isoform c [Homo sapiens] | 3 | 1 | 1 | 1 | 0 |
| NP_009057.1 | 1 | transitional endoplasmic reticulum ATPase isoform 1 [Homo sapiens] | 1 | 1 | 1 | 1 | 0 |
| NP_001341856.1 | 1 | transitional endoplasmic reticulum ATPase isoform 2 [Homo sapiens] | 1 | 1 | 1 | 1 | 0 |
| NP_001273201.1 | 1 | translationally-controlled tumor protein isoform 1 [Homo sapiens] | 7 | 1 | 1 | 1 | 0 |
| NP_003286.1 | 1 | translationally-controlled tumor protein isoform 2 [Homo sapiens] | 8 | 1 | 1 | 1 | 0 |

**Supplementary Table 1: hIPSC-CM secreted Proteins Identified by LC-MS/MS specific to VEEV TC-83 infection.**
| Accession | Biological Replicates (#) | Description | Coverage [%] | Peptides (#) | PSMs (#) | Unique Peptides (#) | Score Sequest HT: A2 Sequest HT |
| --- | --- | --- | --- | --- | --- | --- | --- |
| NP_001273202.1 | 1 | translationally-controlled tumor protein isoform 3 [Homo sapiens] | 9 | 1 | 1 | 1 | 0 |
| NP_001265117.1 | 2 | tropomyosin alpha-3 chain isoform 6 [Homo sapiens] | 8 | 1 | 1 | 1 | 0 |
| NP_001265120.1 | 2 | tropomyosin alpha-3 chain isoform 9 [Homo sapiens] | 9 | 1 | 1 | 1 | 0 |
| NP_079079.1 | 1 | tubulin alpha chain-like 3 isoform 1 [Homo sapiens] | 3 | 1 | 1 | 1 | 0 |
| NP_001165335.1 | 1 | tubulin alpha chain-like 3 isoform 2 [Homo sapiens] | 3 | 1 | 1 | 1 | 0 |
| NP_005991.1 | 1 | tubulin alpha-4A chain isoform 1 [Homo sapiens] | 3 | 1 | 1 | 1 | 0 |
| NP_001265481.1 | 1 | tubulin alpha-4A chain isoform 2 [Homo sapiens] | 3 | 1 | 1 | 1 | 0 |
| XP_016860313.1 | 1 | tubulin alpha-4A chain isoform X1 [Homo sapiens] | 4 | 1 | 1 | 1 | 0 |
| NP_061816.1 | 1 | tubulin alpha-8 chain isoform 1 [Homo sapiens] | 3 | 1 | 1 | 1 | 0 |
| NP_001180343.1 | 1 | tubulin alpha-8 chain isoform 2 [Homo sapiens] | 3 | 1 | 1 | 1 | 0 |
| NP_001345618.1 | 1 | tubulin beta 8 pseudogene 12 [Homo sapiens] | 3 | 1 | 1 | 1 | 0 |
| NP_001276052.1 | 1 | tubulin beta-4A chain isoform 1 [Homo sapiens] | 3 | 1 | 1 | 1 | 0 |
| NP_001276056.1 | 1 | tubulin beta-4A chain isoform 2 [Homo sapiens] | 3 | 1 | 1 | 1 | 0 |
| NP_006078.2 | 1 | tubulin beta-4A chain isoform 3 [Homo sapiens] | 3 | 1 | 1 | 1 | 0 |
| NP_001276060.1 | 1 | tubulin beta-4A chain isoform 4 [Homo sapiens] | 4 | 1 | 1 | 1 | 0 |
| NP_001290453.1 | 1 | tubulin beta-6 chain isoform 1 [Homo sapiens] | 3 | 1 | 1 | 1 | 0 |
| NP_001290455.1 | 1 | tubulin beta-6 chain isoform 3 [Homo sapiens] | 3 | 1 | 1 | 1 | 0 |
| NP_001290456.1 | 1 | tubulin beta-6 chain isoform 4 [Homo sapiens] | 4 | 1 | 1 | 1 | 0 |
| NP_001290457.1 | 1 | tubulin beta-6 chain isoform 5 [Homo sapiens] | 4 | 1 | 1 | 1 | 0 |
| NP_001290458.1 | 1 | tubulin beta-6 chain isoform 6 [Homo sapiens] | 5 | 1 | 1 | 1 | 0 |
| NP_817124.1 | 1 | tubulin beta-8 chain [Homo sapiens] | 3 | 1 | 1 | 1 | 0 |
| XP_016871681.1 | 1 | tubulin beta-8 chain isoform X1 [Homo sapiens] | 4 | 1 | 1 | 1 | 0 |
| XP_011517762.1 | 1 | tubulin beta-8 chain isoform X2 [Homo sapiens] | 5 | 1 | 1 | 1 | 0 |
| XP_011517761.1 | 1 | tubulin beta-8 chain isoform X3 [Homo sapiens] | 4 | 1 | 1 | 1 | 0 |
| XP_016883529.1 | 1 | 14-3-3 protein beta/alpha isoform X2 [Homo sapiens] | 13 | 1 | 1 | 1 | 0 |
| XP_016883841.1 | 1 | interferon-induced GTP-binding protein Mx1 isoform X2 [Homo sapiens] | 2 | 1 | 1 | 1 | 0 |
| XP_016884902.1 | 2 | glypican-3 isoform X1 [Homo sapiens] | 3 | 1 | 1 | 1 | 0 |
| VEEV_E2 | 1 | VEEV_E2 | 6 | 1 | 1 | 1 | 0 |
| XP_024305231.1 | 1 | protein AHNAK2 isoform X1 [Homo sapiens] | 0 | 1 | 1 | 1 | 0 |
| XP_024306911.1 | 1 | tubulin beta 8 pseudogene 12 isoform X1 [Homo sapiens] | 4 | 1 | 1 | 1 | 0 |
| XP_024306913.1 | 1 | tubulin beta 8 pseudogene 12 isoform X2 [Homo sapiens] | 4 | 1 | 1 | 1 | 0 |
| XP_024306914.1 | 1 | tubulin beta 8 pseudogene 12 isoform X3 [Homo sapiens] | 4 | 1 | 1 | 1 | 0 |
| XP_024306915.1 | 1 | tubulin beta 8 pseudogene 12 isoform X4 [Homo sapiens] | 4 | 1 | 1 | 1 | 0 |
| XP_024306916.1 | 1 | tubulin beta 8 pseudogene 12 isoform X5 [Homo sapiens] | 5 | 1 | 1 | 1 | 0 |
| XP_024307843.1 | 1 | amyloid-beta A4 protein isoform X1 [Homo sapiens] | 2 | 1 | 1 | 1 | 0 |

